# DeepMRG: a multi-label deep learning classifier for predicting bacterial metal resistance genes

**DOI:** 10.1101/2023.11.14.566903

**Authors:** Muhit Islam Emon, Liqing Zhang

**Author notes:** Corresponding author; (LZ). (MIE).

## Abstract

The widespread misuse of antibiotics has escalated antibiotic resistance into a critical global public health concern. Beyond antibiotics, metals function as antibacterial agents. Metal resistance genes (MRGs) enable bacteria to tolerate metal-based antibacterials and may also foster antibiotic resistance within bacterial communities through co-selection. Thus, predicting bacterial MRGs is vital for elucidating their involvement in antibiotic resistance and metal tolerance mechanisms. The “best hit” approach is mainly utilized to identify and annotate MRGs. This method is sensitive to cutoff values and produces a high false negative rate. Other than the best hit approach, only a few antimicrobial resistance (AMR) detection tools exist for predicting MRGs. However, these tools lack comprehensive annotation for MRGs conferring resistance to multiple metals. To address such limitations, we introduce DeepMRG, a deep learning-based multi-label classifier, to predict bacterial MRGs. Because a bacterial MRG can confer resistance to multiple metals, DeepMRG is designed as a multi-label classifier capable of predicting multiple metal labels associated with an MRG. It leverages bit score-based similarity distribution of sequences with experimentally verified MRGs. To ensure unbiased model evaluation, we employed a clustering method to partition our dataset into six subsets, five for cross-validation and one for testing, with non-homologous sequences, mitigating the impact of sequence homology. DeepMRG consistently achieved high overall F1-scores and significantly reduced false negative rates across a wide range of datasets. It can be used to predict bacterial MRGs in metagenomic or isolate assemblies. The web server of DeepMRG can be accessed at https://deepmrg.cs.vt.edu/deepmrg and the source code is available at https://github.com/muhit-emon/DeepMRG under the MIT license.

## Introduction

Antibiotic resistance poses a significant threat to global human health and is increasingly becoming a silent pandemic due to the widespread and inappropriate use of antibiotics [1, 2]. In response, there has been an increased reliance on metal-based antibacterial agents. However, it is important to note that bacteria can develop resistance to metals through exposure [3]. Moreover, the use of these antibacterial metals can contribute to the emergence and persistence of antibiotic resistance in bacterial populations through co-selection [4, 5]. Therefore, it is crucial to systematically and comprehensively detect and annotate bacterial metal resistance genes (MRGs) to gain insights into their role in developing antibiotic resistance and to understand the key mechanisms behind bacterial tolerance to metals.

The prediction of an MRG is primarily conducted using the “best hit” method, which involves comparing the query gene sequence to existing reference databases using programs such as BLAST [6] and DIAMOND [7] and annotating the gene’s function based on the reference sequence it shows the highest similarity to [8, 9]. However, the best hit method requires setting identity cutoff scores (and/or alignment lengths) and is sensitive to these cutoff values, making it challenging to decide on an appropriate threshold. Generally, a high identity cutoff is applied when using the best hit method to predict bacterial MRGs. For instance, the authors in [10] and [11] employed the best hit method with an identity greater than 80% to reference sequences to predict MRGs. While the best hit method with a high cutoff value generally exhibits a low false positive rate [12], it can result in a high false negative rate [8, 13].

Apart from the best-hit approach, MEGARes 3.0 (AMR++ 3.0) [14], AMR-meta [15], and AMRFinderPlus [16] can identify MRGs. MEGARes 3.0 (AMR++ 3.0) and AMR-meta are tailored for metagenomics short-reads and are not suitable for predicting MRGs in contigs. Additionally, These programs lack detailed annotation for MRGs that confer resistance to multiple metals. For instance, genes associated with resistance to multiple metals are annotated in these programs to a general class called multi-metal resistance, and no details are provided as to what kinds of metals are included in “multi-metal”. On the other hand, AMRFinderPlus can detect MRGs from protein sequences and provide detailed annotations for MRGs conferring resistance to multiple metals using BLAST and HMMER searches. However, its annotation is limited to a subset of possible multi-label scenarios, as per the BacMet [5] databases, the most comprehensive databases of bacterial metal resistance genes.

Here, we introduce DeepMRG, a multi-label classifier that utilizes deep learning to predict bacterial MRGs. We designed DeepMRG as a multi-label classifier since an MRG can confer resistance to multiple metals. Our model can provide specific annotations for multi-metal resistance genes, indicating the particular metals to which the gene confers resistance. DeepMRG aligns a query gene sequence with experimentally confirmed MRGs, extracting alignment bit scores. These alignment bit scores are then used to derive the similarity distribution of the query sequence with 66 types of experimentally verified MRGs. This bit score-based similarity distribution serves as the feature for the deep neural networks. To minimize the effect of sequence homology on model evaluation, we created the training, validation, and test datasets with a clustered split method. DeepMRG demonstrated good predictive performance for MRGs during both 5-fold cross-validation and on the test dataset. Furthermore, we assessed DeepMRG’s ability to identify and classify sequences with low similarity to experimentally confirmed MRGs. DeepMRG was also validated using an independent set of heavy metal resistance genes and *in silico* spike-in experiment. DeepMRG excelled in precision, recall, and F1-score in all the conducted experiments, with notably lower false negative rates. DeepMRG is implemented as an easy-to-use web server available via https://deepmrg.cs.vt.edu/deepmrg and as a command line tool freely available at https://github.com/muhit-emon/DeepMRG. It is fully documented in S1 File.

## Materials and methods

### Data collection and processing

We collected antibacterial biocide and metal resistance genes from BacMet [5]. BacMet contains 753 gene sequences in a database named BacMet EXP DB where the genes have been experimentally verified to confer resistance to metals and/or antibacterial biocides. Additionally, it provides BacMet Predicted DB, a database containing 155,512 potential resistance genes compiled from public sequence repositories based on sequence homology to genes with experimentally verified resistance functions.

We focus on predicting bacterial MRGs in this paper. Therefore, we extracted MRGs from BacMet databases by searching for metal names in gene metadata, excluding genes that confer resistance only to antibacterial biocides. It left us 485 sequences in BacMet EXP MRG DB and 93,367 sequences in BacMet Predicted MRG DB, as shown in Fig 1A. BacMet EXP MRG is used throughout the paper to denote a gene in BacMet EXP MRG DB, and BacMet Predicted MRG is used to indicate a gene sequence belonging to BacMet Predicted MRG DB.

**Fig 1.**
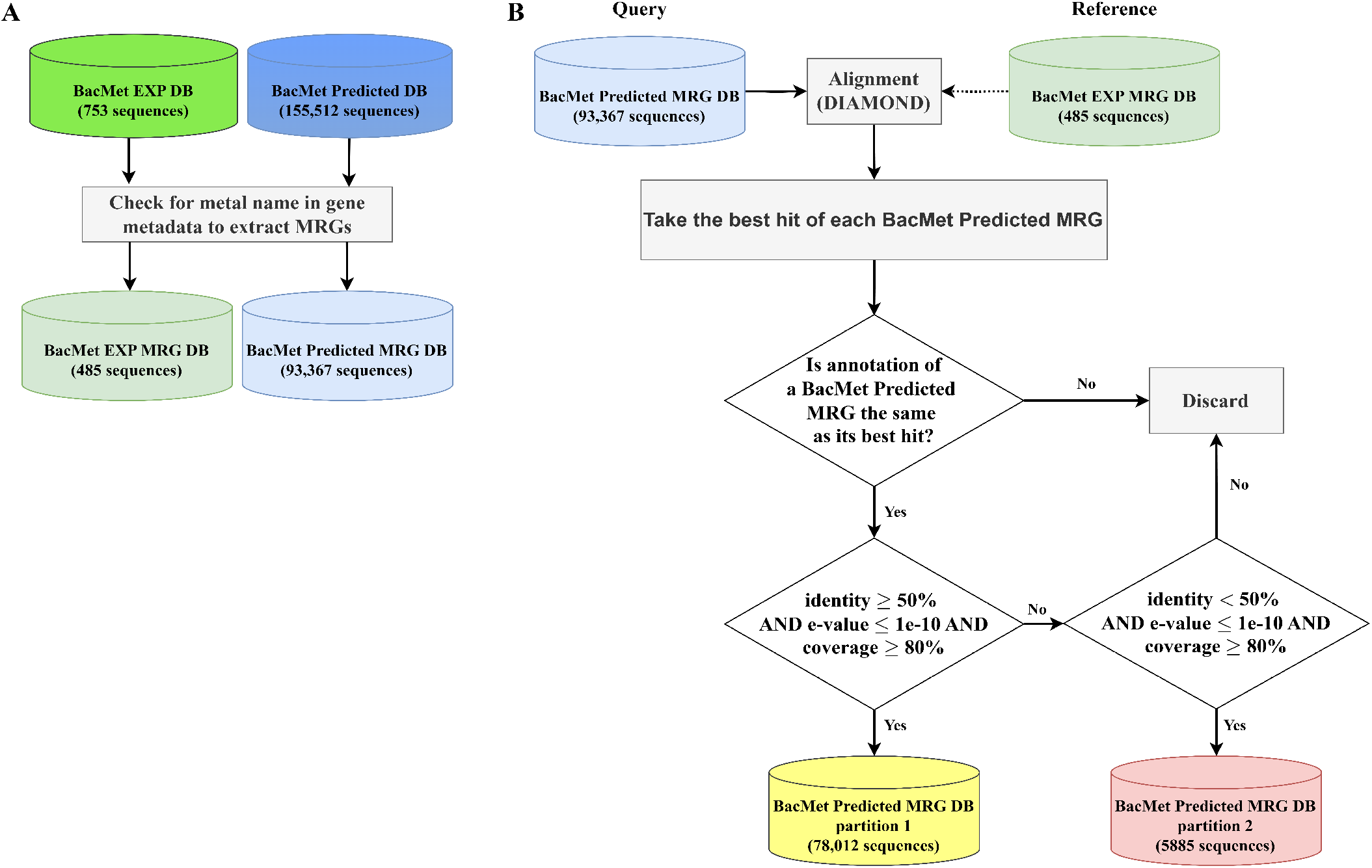
Collection and processing of data. A: Extraction of MRGs from BacMet [5] databases by searching metal names in gene metadata. B: Partition of BacMet Predicted MRG DB sequences into two databases. BacMet Predicted MRGs were aligned against the experimentally verified MRGs using DIAMOND [7]. The best hit was selected for each BacMet Predicted MRG and a set of filters were applied to create BacMet Predicted MRG DB partitions 1 and 2.

We validated the annotations of BacMet Predicted MRGs by taking their best hit to BacMet EXP MRG DB. We employed DIAMOND [7], a tool similar to BLAST [6] but considerably faster, to align the BacMet Predicted MRGs against BacMet EXP MRG DB and identified their best hit. Based on the sequence identity, e-value, and coverage of the best hit alignment, the BacMet Predicted MRGs were divided into the following two databases, as shown in Fig 1B:

1. **BacMet Predicted MRG DB partition 1**: A BacMet Predicted MRG is included in the BacMet Predicted MRG DB partition 1 if its best hit to a BacMet EXP MRG has ≥50% sequence identity, e-value ≤ 1e-10, alignment coverage ≥ 80%, and it has the same metal resistance annotation to the BacMet EXP MRG. 78,012 genes passed these constraints and were added to the BacMet Predicted MRG DB partition 1. We set the cutoffs of identity and e-value following [17] where these thresholds were used to identify high and mid quality antibiotic resistance genes from public databases. The alignment coverage cutoff was selected following [5].
2. **BacMet Predicted MRG DB partition 2**: A BacMet Predicted MRG is added to the BacMet Predicted MRG DB partition 2 if its best hit to a BacMet EXP MRG has *<* 50% sequence identity, e-value ≤ 1e-10, alignment coverage ≥ 80%, and possesses identical metal resistance annotation as the BacMet EXP MRG. After satisfying all these constraints, 5885 genes were placed in the BacMet Predicted MRG DB partition 2.

We used the sequences in BacMet Predicted MRG DB partition 1 to construct the training, validation, and test datasets for our DeepMRG model. We held out the sequences in BacMet Predicted MRG DB partition 2 and the test set sequences for evaluating and comparing the performance of DeepMRG with the BLAST best hit method and AMRFinderPlus [16].

### Construction of training, validation, and test datasets using clustered split

We created the training, validation, and test datasets from the BacMet Predicted MRG DB partition 1 following the non-homologous database split technique employed in [18]. The sequences in BacMet Predicted MRG DB partition 1 are categorized into 63 different types based on the metal labels they confer resistance to (types and the number of sequences within each type are presented in S2 File). We utilized MMseqs2 [19] to cluster the sequences in each type using 40% sequence identity and 50% coverage thresholds. Then, the clusters in each type were randomly split into six sets, where an entire cluster was placed in one of the six sets. Subsequently, we combined the corresponding sets from all 63 types and obtained six datasets (D1-D5 and TEST). Datasets D1 to D5 were used for 5-fold cross-validation, and the dataset TEST was for testing, as shown in Fig 2. Constructing training, validation, and test datasets utilizing this clustered split approach reduces the impact of sequence homology on evaluating the deep learning model.

**Fig 2.**
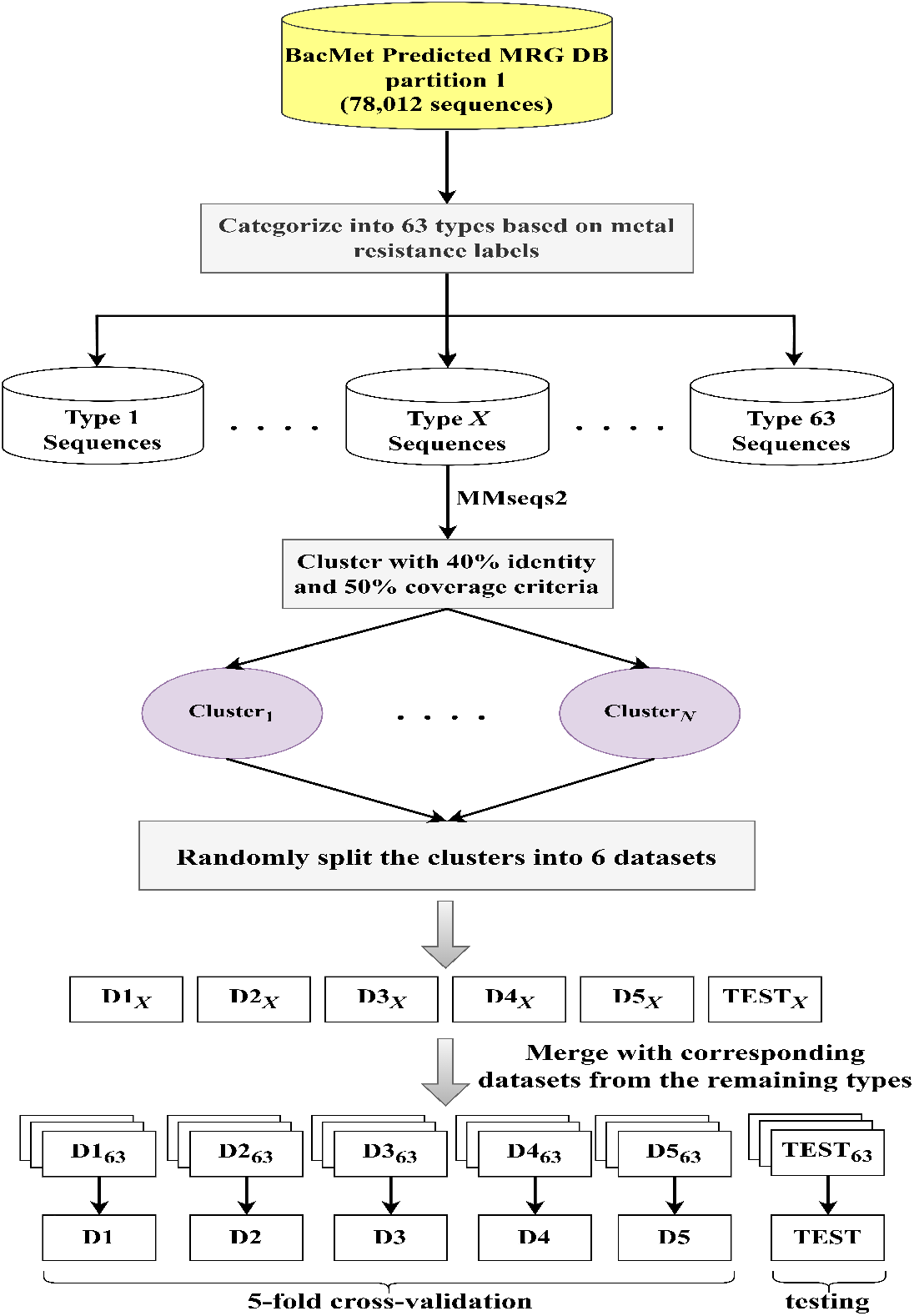
Clustered split. The sequences in BacMet Predicted MRG DB partition 1 are categorized into 63 types based on their metal resistance labels. The sequences in each type (type *X* in this figure where 1 ≤ *X* ≤ 63) were grouped into clusters using MMseqs2 [19] at 40% identity and 50% coverage. The clusters were randomly partitioned in six sets (D1_*X*_-D5_*X*_ and TEST_*X*_) where an entire cluster was included in one of the six sets. Finally, the corresponding datasets from the remaining types were combined to make the datasets used for 5-fold cross-validation and testing (D1-D5 and TEST) of our deep learning model.

### Feature extraction

Deep learning models require a sequence to be represented as a vector of numerical values called features for prediction or classification tasks. In this paper, we adopted the concept of bit score-based similarity distribution used in [17] and [20]. The features are the alignment bit scores between full-length gene sequences and 66 types of experimentally verified MRGs available in BacMet EXP MRG DB. We used the bit score as the indicator of sequence similarity because it is independent of the database size, unlike the e-value [21]. The process for computing features for a full-length gene sequence is outlined as follows (refer to Fig 3):

**Fig 3.**
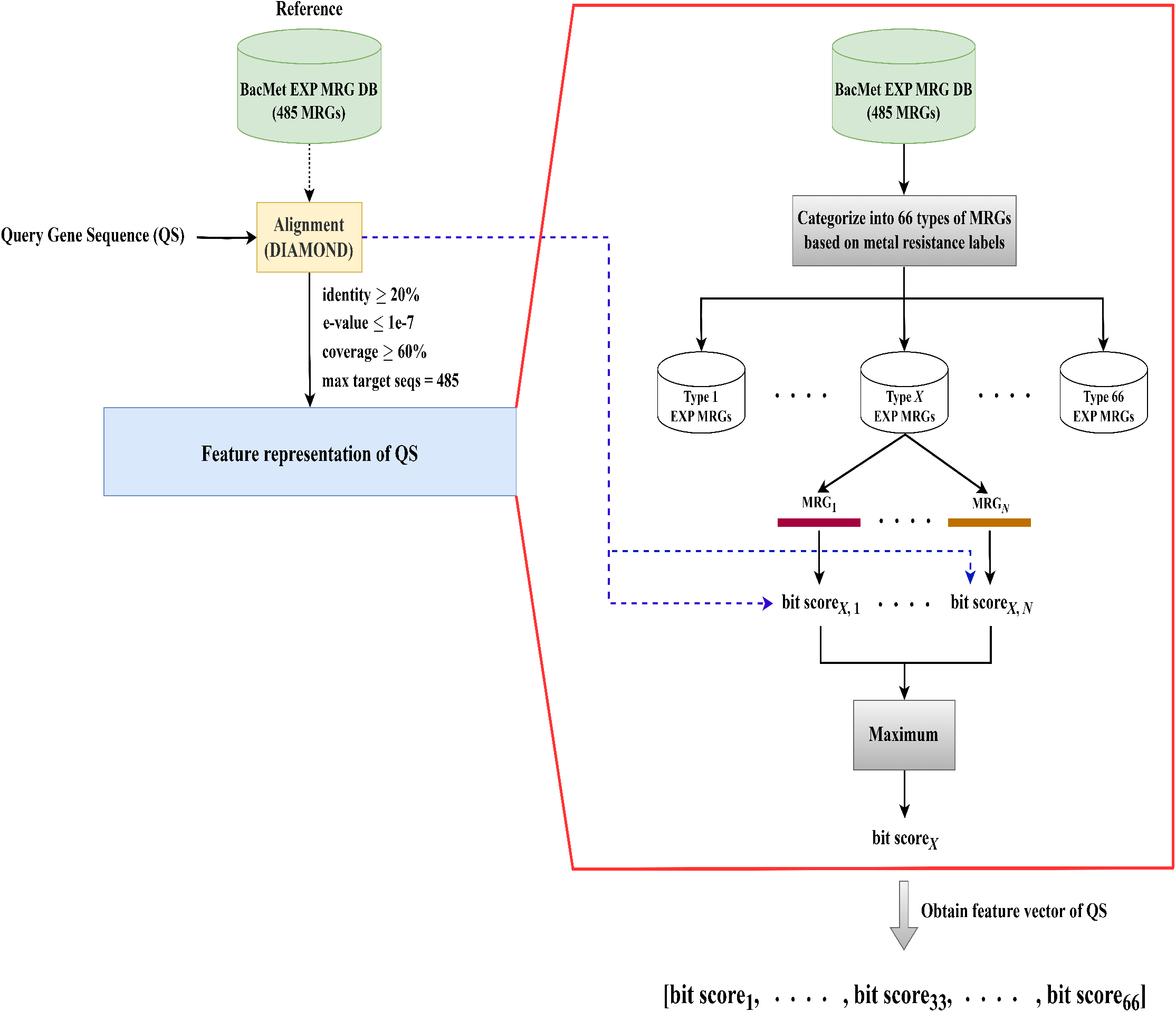
The feature vector construction process of a query gene sequence (QS). QS is aligned with the experimentally verified bacterial MRGs using DIAMOND [7]. BacMet EXP MRGs are categorized into 66 types according to their metal resistance labels. We use type *X* (1 ≤ *X* ≤ 66) in this figure to show the feature extraction process for QS. We assume that *N* experimentally verified MRGs are in type *X*. We get the bit scores between QS and these *N* MRGs from the alignment step. Selecting the highest bit score among these *N* scores yields bit score_*X*_, representing the similarity between QS and type *X* of BacMet EXP MRGs. Similarly, we get 66 such bit scores for each of the 66 types. The feature vector *V*_*QS*_ ∈ ℝ^66^ of QS contains these bit scores as feature values.

1. The query gene sequence (QS) is aligned to the experimentally confirmed 485 MRGs in BacMet EXP MRG DB using DIAMOND under the ‘very sensitive’ parameter with the alignment constraints: a minimum sequence identity of 20%, an e-value lower than 1e-7, and a minimum alignment coverage of 60%. If QS does not have alignment with any of the 485 BacMet EXP MRGs, it is filtered out and not considered for further prediction. Otherwise, we go to the following steps to calculate the feature vector of QS. The alignment step works as a filter and only passes MRG-like sequences to our deep learning model for prediction.
2. BacMet EXP MRGs are categorized into 66 types according to their metal resistance labels (types and the number of sequences within each type are presented in S3 File).
3. Here, we use the type *X* (1 ≤ *X* ≤ 66) to demonstrate how we compute the feature vector for QS. Assume that there are *N* MRGs in type *X*. The alignment bit scores between QS and these *N* MRGs are obtained from step 1. Subsequently, we take the maximum among these *N* bit scores, yielding bit score_*X*_, which serves as the representative bit score between QS and type *X* of experimentally verified MRGs.

Similarly to the previous procedure, we get 66 bit scores which are then normalized to the [0, 1] interval to represent the similarity distribution of QS with 66 types of experimentally confirmed bacterial MRGs. Thus, the feature vector of QS for DeepMRG is *V*_*QS*_ ∈ ℝ^66^ where *V*_*QS*_ = [bit score_1_, …, bit score_33_, …, bit score_66_].

### Deep learning model architecture and training

Subsequently, we trained DeepMRG, a multi-label deep neural network model, for classifying a gene sequence into one or multiple metal resistance categories by taking into account the similarity distribution of the sequence to 66 types of experimentally verified bacterial MRGs. Deep learning models can discern relevant features without human interference, which is one of its key advantages [22]. The DeepMRG model consists of an input layer, three hidden dense layers with 55, 45, and 35 nodes respectively, and an output layer. Each hidden dense layer utilizes the ELU activation function. To prevent overfitting, we applied dropout regularization with a dropout rate of 10% after each hidden layer. The Adam optimizer was used to update the weights, and binary cross entropy served as the loss function. The output layer consists of 23 nodes (one per metal). We employed the sigmoid activation function in the output layer, which computes the likelihood scores for each metal category. The model architecture is shown in Fig 4. The model was implemented using Python 3.6 with Scikit-learn 0.24.2 and Keras 2.2.0 with Tensorflow 2.2.0 backend.

**Fig 4.**
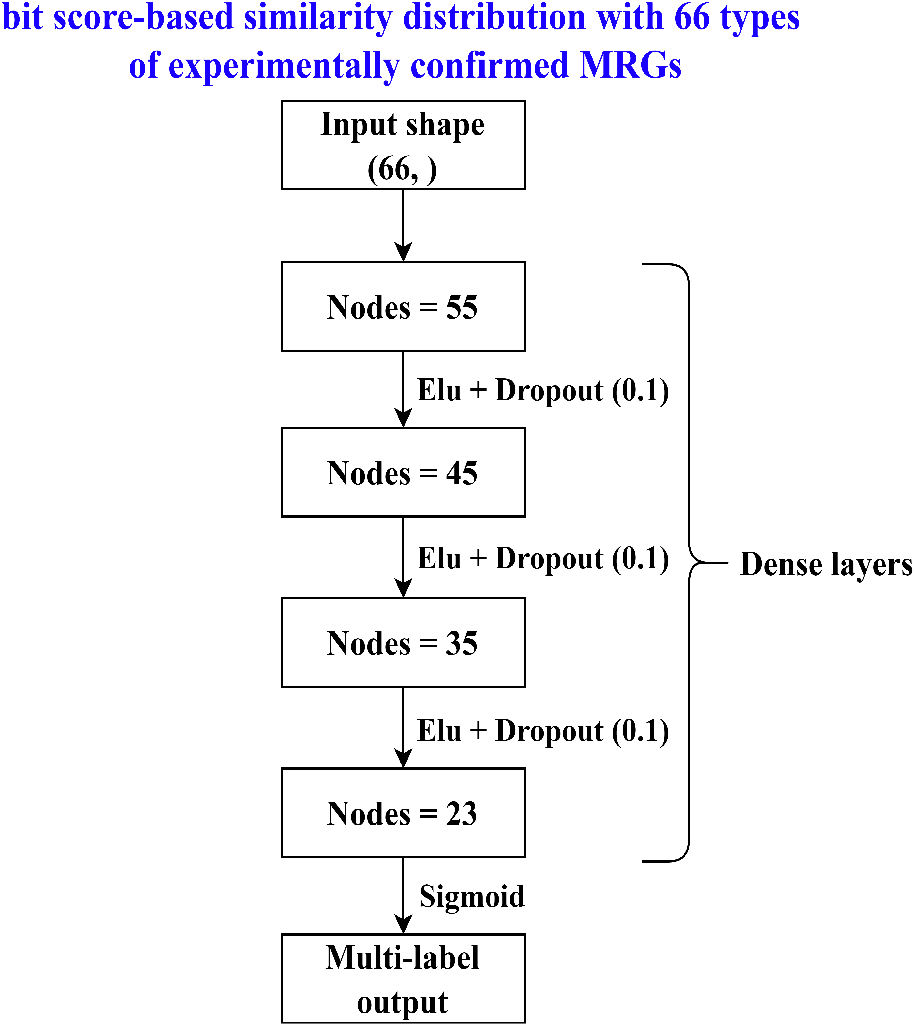
DeepMRG model architecture.

We performed 5-fold cross-validation using sets D1 to D5 (see Fig 2), training five deep neural networks, each sequentially using one set for validation and the remaining four for training.

In 5-fold cross-validation, we trained five deep neural networks. For an input query gene sequence, these five neural networks produce likelihood scores for each metal category. Then, we sum the scores from the five neural networks for each metal (the maximum achievable score is 5 per metal). The query sequence is then classified into the metal categories with the corresponding score of more than a threshold (default 3.5).

### Competing methods

We compared our tool with the BLAST [6] best hit method and AMRFinderPlus [16] for classifying MRGs using protein or gene sequences as input.

#### BLAST best hit

BLAST [6] is one of the widely used sequence alignment tools. We aligned the query sequences against the BacMet experimentally confirmed MRG database using BLASTp and took the best hit to annotate them. We ran the BLASTp program with the option ‘-max target seqs 1’ and utilized various sequence identity cutoffs as the representatives of the best hit approach.

#### AMRFinderPlus

AMRFinderPlus [16] classifies MRGs by employing BLAST and HMMER searches against its reference gene catalog. This catalog encompasses antimicrobial resistance, acid, biocide, metal, heat resistance genes, and virulence genes. We ran the AMRFinderPlus program with the default parameters and utilized the ‘plus’ subset of the reference gene catalog (database version 2024-1-31.1), which includes genes associated with metal resistance.

## Results

### Performance evaluation metrics

We evaluated the performance of DeepMRG using both label-based and sample-based metrics. Label-based metrics assess the model’s performance based on the prediction of class labels. We used precision, recall, and F1-score as label-based metrics. We also reported both macro-average and weighted-average results. The macro-average calculates the average performance of all classes, treating each class equally. On the other hand, the weighted average takes into account the number of sequences in each class, providing a performance measure that is influenced by the class distribution. The weighted-average is particularly useful when the class sizes are imbalanced.

Additionally, we utilized sample-based metrics to evaluate the performance of our model. Sample-based metrics assess the model’s performance on an individual sample level within the dataset. As sample-based metrics, we reported samples-average precision, samples-average recall, and samples-average F1-score. A detailed description of the performance evaluation metrics is provided in S4 File.

### Performance of DeepMRG under 5-fold cross-validation

We evaluated the performance of DeepMRG through 5-fold cross-validation using clustered splits (see Fig 2). The overall F1-scores, including their mean and standard deviation, obtained from all five cross-validation experiments, are presented in Table 1. DeepMRG consistently achieved high F1-scores, with an average macro-average of 98.2%, an average weighted-average of 98.8%, and an average samples-average of 98.4%. These results indicate that DeepMRG can accurately predict MRGs and exhibits good generalization capabilities as a deep learning model. Furthermore, the standard deviations of 1.3% for macro-average F1-scores, 0.4% for weighted-average, and 0.5% for samples-average F1-scores across five independent cross-validation experiments highlight DeepMRG’s stability. Detailed classification reports for each metal, including precision, recall, and F1-score per fold, can be found in S5 File.

**Table 1.** The 5-fold cross-validation results of DeepMRG to predict MRGs.

| Fold | Macro-average F1-score | Weighted-average F1-score | Samples-average F1-score |
| --- | --- | --- | --- |
| 1st | 0.99 | 0.99 | 0.99 |
| 2nd | 0.99 | 0.99 | 0.98 |
| 3rd | 0.98 | 0.98 | 0.98 |
| 4th | 0.99 | 0.99 | 0.98 |
| 5th | 0.96 | 0.99 | 0.99 |
| Mean ± SD | 0.982 ± 0.013 | 0.988 ± 0.004 | 0.984 ± 0.005 |

Performance assessment of DeepMRG through 5-fold cross-validation. Here, SD stands for standard deviation.

### Performance on the test dataset

We compared DeepMRG with AMRFinderPlus and BLAST best hit approaches on the test dataset comprising 11,447 sequences created using a clustered split method, as shown in Fig 2. The overall performance of these methods on the test dataset is presented in Table 2, and the individual metal category-wise results are shown in Fig 5. Upon analyzing the reference gene catalog of AMRFinderPlus, we found that it identifies MRGs associated with resistance to 12 metals, including arsenic (As), cadmium (Cd), chromium (Cr), cobalt (Co), copper (Cu), gold (Au), lead (Pb), mercury (Hg), nickel (Ni), silver (Ag), tellurium (Te), and zinc (Zn). However, on our test dataset, AMRFinderPlus could not detect any MRGs conferring resistance to cadmium (Cd), cobalt (Co), lead (Pb), tellurium (Te), and zinc (Zn). Additionally, it exhibited poor recall rates for arsenic (As), chromium (Cr), copper (Cu), nickel (Ni), and silver (Ag), while achieving very high F1-scores for gold (Au) and mercury (Hg). BLAST best hit with an identity cutoff at 80% (BLAST-80%), a common choice in prior studies [10, 11] to predict bacterial MRGs, achieved very high precision (≥ 0.99) for all metal categories on our test dataset. However, its recall was compromised for many metals, impacting the overall F1-scores. BLAST best hit with an identity cutoff of 60% (BLAST-60%) maintained the same precision levels as BLAST-80% while significantly improving recall across all metal categories, resulting in a higher overall F1-scores on the test dataset. DeepMRG also exhibited very high precision (≥ 0.98) for all metal labels, akin to BLAST-80% and BLAST-60%. It demonstrated similar or better recall rates than the best hit methods across all metals except copper (Cu) and achieved better F1-scores than AMRFinderPlus for most metal categories. For gold (Au) and mercury (Hg) resistance, both DeepMRG and AMRFinderPlus performed equally well. Overall, DeepMRG outperformed AMRFinderPlus and BLAST-80% in terms of F1-scores. It achieved a similar weighted-average F1-score and samples-average F1-score as BLAST-60%, along with a slightly better macro-average F1-score than BLAST-60%.

**Table 2.** Performance on test dataset.

| Methods | Macro-average F1-score | Weighted-average F1-score | Samples-average F1-score |
| --- | --- | --- | --- |
| DeepMRG | 0.99 | 0.98 | 0.95 |
| AMRFinderPlus | 0.31 | 0.17 | 0.10 |
| BLAST best hit (80%, 1e-7, 60%) | 0.87 | 0.84 | 0.71 |
| BLAST best hit (60%, 1e-7, 60%) | 0.98 | 0.98 | 0.95 |

Performance of DeepMRG, AMRFinderPlus, and BLAST best hit methods under different parameters on test dataset. Here, BLAST best hit (x%, y, z%) refers to the best alignment hit with identity ≥ x%, e-value ≤ y, and coverage ≥ z%.

**Fig 5.**
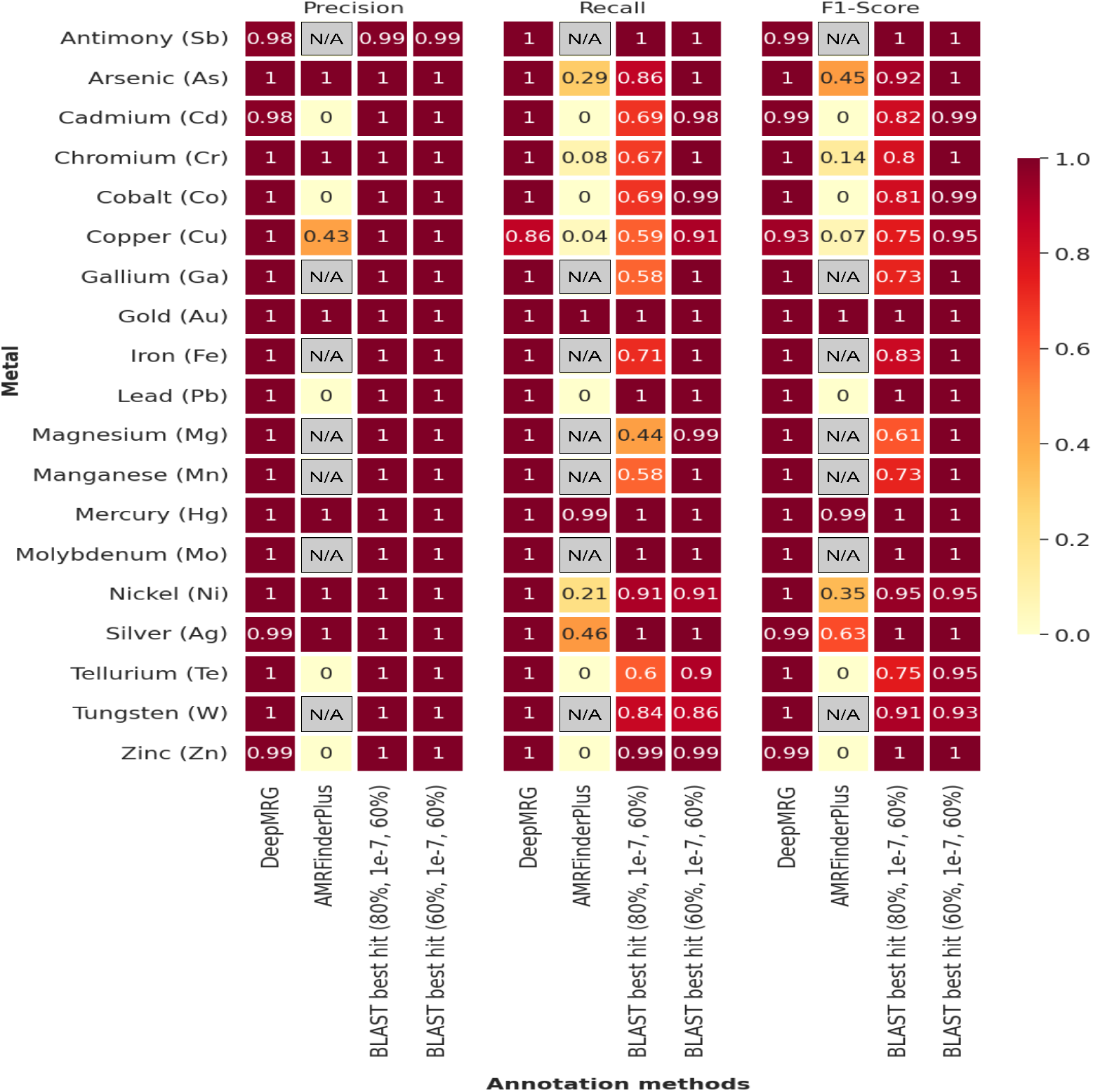
Precision, recall, and F1-score comparison among DeepMRG, AMRFinderPlus, and BLAST best hit approaches with different parameter settings for individual metal categories in the test dataset. The precision, recall, and F1-score values for metal categories where AMRFinderPlus does not provide predictions are denoted as N/A. BLAST best hit (x%, y, z%) in this figure refers to the best alignment hit with identity ≥ x%, e-value ≤ y, and coverage ≥ z%. As a result of the clustered split, not all 23 metal labels are present in the test dataset.

### Performance on the sequences in BacMet Predicted MRG DB partition 2

We employed DeepMRG to classify the 5885 sequences from BacMet Predicted MRG DB Partition 2 and conducted a performance comparison with AMRFinderPlus and BLAST best hit approaches. The construction process of this database is detailed earlier in the “data collection and processing” section. None of these 5885 sequences were utilized for training DeepMRG. Notably, 75% of these sequences exhibit less than 60% sequence identity with our training set, as shown in S1 Fig. Because the sequences in BacMet Predicted MRG DB Partition 2 have less than 50% sequence identity with the BacMet EXP MRGs (see Fig 1), using a 50% or greater identity cutoff in the BLAST best hit method would result in these potential MRGs not being detected. Therefore, in this section, we employed 40% and 30% identity cutoff values for the BLAST best hit method (namely BLAST-40% and BLAST-30%) to compare with DeepMRG. BLAST-40% achieved similar or better precision compared to BLAST-30% on individual metal labels, while BLAST-30% exhibited higher recall rates (see S6 File). The overall F1-scores of DeepMRG, AMRFinderPlus, BLAST-40%, and BLAST-30% are presented in Table 3. AMRFinderPlus performed poorly on the BacMet Predicted MRG DB partition 2 dataset, only identifying MRGs associated with resistance to mercury (Hg) with an F1-score of 0.5. DeepMRG outperformed AMRFinderPlus, BLAST-40%, and achieved overall F1-scores highly comparable to BLAST-30%. For most metal categories, DeepMRG demonstrated higher F1-scores than BLAST-40% and F1-scores on par with BLAST-30%, except for silver (Ag).

**Table 3.**
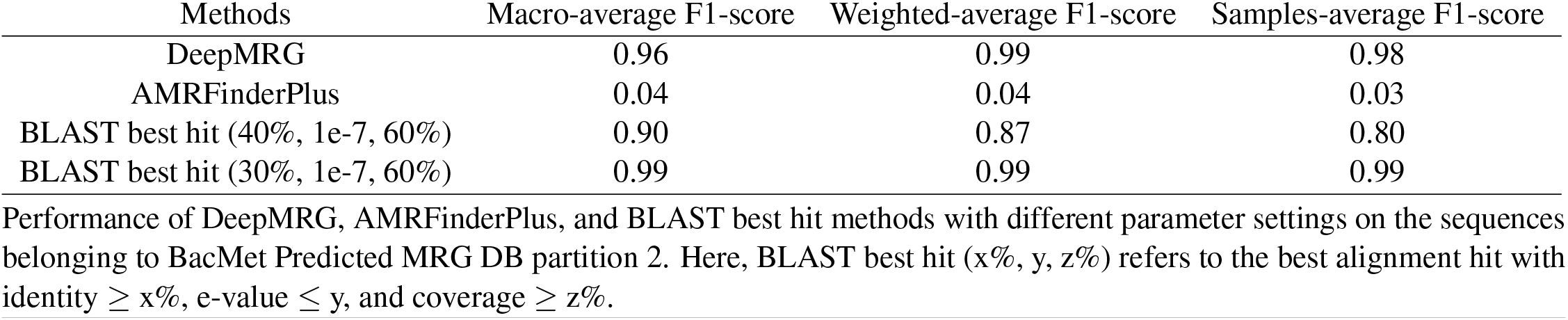
Performance on BacMet Predicted MRG DB partition 2.

| Methods | Macro-average F1-score | Weighted-average F1-score | Samples-average F1-score |
| --- | --- | --- | --- |
| DeepMRG | 0.96 | 0.99 | 0.98 |
| AMRFinderPlus | 0.04 | 0.04 | 0.03 |
| BLAST best hit (40%, 1e-7, 60%) | 0.90 | 0.87 | 0.80 |
| BLAST best hit (30%, 1e-7, 60%) | 0.99 | 0.99 | 0.99 |

Performance of DeepMRG, AMRFinderPlus, and BLAST best hit methods with different parameter settings on the sequences belonging to BacMet Predicted MRG DB partition 2. Here, BLAST best hit (x%, y, z%) refers to the best alignment hit with identity ≥ x%, e-value ≤ y, and coverage ≥ z%.

This comparison underscores the best hit approach’s sensitivity to sequence identity cutoffs, while DeepMRG eliminates the need for these threshold settings, delivering consistent and excellent performance in predicting MRGs across diverse datasets.

### Validation through an independent set of heavy metal resistance genes

Based on an independent study by Klonowska *et al*. [23], we obtained 53 heavy metal resistance (HMR) genes from heavy metal-tolerant *Cupriavidus* strain STM 6070 to evaluate DeepMRG’s ability to predict novel bacterial MRGs. In this study, the authors conducted wet lab experiments to demonstrate that STM 6070 exhibits significantly higher tolerance to Ni^2+^and Zn^2+^concentrations compared to *Cupriavidus taiwanensis* strains. Moreover, computational and comparative genomics approaches were used to identify HMR genes in the STM 6070 genome, potentially involved in arsenic, cadmium, chromium, cobalt, copper, nickel, silver, and zinc resistance.

Employing a sequence identity threshold of 50% or more in the best hit approach is a common practice in bioinformatics [24]. Therefore, we set the identity cutoff parameter to 50% for the BLAST best hit method (BLAST-50%) and compared its results with those of DeepMRG for the independent dataset. AMRFinderPlus failed to detect any HMR gene in the independent dataset using its default parameters. Subsequently, we re-ran the tool with the identity parameter set to 50% using the option ‘-i 0.5’, denoted as AMRFinderPlus-50%. The prediction results of DeepMRG, AMRFinderPlus-50%, and BLAST-50% in terms of F1-scores for the HMR genes in the independent dataset are shown in Table 4. BLAST best hits of these HMR genes against the experimentally validated MRGs, along with details on sequence identity and alignment coverage, are presented in S7 File. While BLAST-50% exhibited better precision for nickel (Ni) and silver (Ag), DeepMRG consistently achieved equal or higher recall than BLAST-50% for all eight metal labels in the independent set (refer to Table 1 of S8 File). AMRFinderPlus-50%, on the other hand, was unable to identify any HMR genes associated with resistance to cadmium (Cd), chromium (Cr), cobalt (Co), and zinc (Zn). Although it showed better precision than DeepMRG for predicting nickel (Ni) resistance, DeepMRG demonstrated higher F1-scores than AMRFinderPlus-50% for all eight metals available in the independent dataset, surpassing it in overall F1-scores. Additionally, DeepMRG yielded higher F1-scores than BLAST-50% for all metals except nickel (Ni), outperforming BLAST-50% in overall F1-scores. It is noteworthy that STM 6070 displays higher tolerance to Ni^2+^and Zn^2+^, and DeepMRG achieved high recall (≥ 0.75) for both metals.

**Table 4.** Performance on the independent dataset.

| Methods | Macro-average F1-score | Weighted-average F1-score | Samples-average F1-score |
| --- | --- | --- | --- |
| DeepMRG | 0.72 | 0.75 | 0.71 |
| AMRFinderPlus-50% | 0.20 | 0.14 | 0.17 |
| BLAST best hit (50%, 1e-7, 60%) | 0.58 | 0.56 | 0.47 |

Performance of DeepMRG, AMRFinderPlus with 50% identity threshold (AMRFinderPlus-50%), and BLAST best hit method at 50% identity cutoff (BLAST-50%) on the independent set of *Cupriavidus* strain STM 6070 HMR genes. BLAST best hit (x%, y, z%) refers to the best alignment hit with identity ≥ x%, e-value ≤ y, and coverage ≥ z%.

We found that out of these 53 HMR genes, six were initially filtered out by our alignment step due to high e-value and low coverage. Adjusting the e-value and coverage cutoffs to 1e-3 and 40%, respectively, following [5, 25], we re-evaluated DeepMRG on the independent dataset. Although two genes were still filtered out during the alignment step, DeepMRG demonstrated improved performance as shown in Tables 2 and 3 of S8 File.

### Validation of DeepMRG through an *in silico* spike-in experiment

MRGs might constitute only a minor portion of the genes in real-world microbial datasets. Thus, assessing how well DeepMRG performs when non-target genes are prevalent is essential. To evaluate DeepMRG’s ability to predict a small number of MRGs within a majority of non-target genes, we constructed a negative microbial dataset simulating a spike-in metagenomic experiment. In this section, our primary emphasis is on the binary classification performance of DeepMRG, specifically in distinguishing positive samples (MRGs) from negative samples. To create negative samples, we collected 27,036 bacterial housekeeping genes from UniProt [26], using 184 different Gene Ontology (GO) terms obtained from [27]. The associated GO terms and biological functions for these genes can be found in S9 File. Employing MMseqs2 at 30% identity and 50% coverage, we conducted clustering of the housekeeping genes together with the experimentally validated MRGs. Those housekeeping genes sharing a cluster with an experimentally confirmed MRG were excluded, resulting in a final set of 26,377 genes as negative samples. Next, we selected the 53 HMR genes from the independent dataset to serve as positive samples. The resulting spike-in dataset comprised a total of 26,430 genes, with the positive samples accounting for (53*/*26, 430)% = 0.2%. DeepMRG achieved an 85% recall rate, identifying 45 out of 53 HMR genes, with a very low false positive rate (*<* 1%). This highlights DeepMRG’s efficacy in predicting MRGs within a large pool of negative samples, mirroring real-world scenarios where MRGs account for a small fraction of the total genes.

### Application of DeepMRG on metagenomic or isolate assembly data

DeepMRG can be applied to metagenomic or isolate assemblies to predict bacterial MRGs. We implemented a pipeline, as shown in Fig 6, for predicting bacterial MRGs from metagenomic or isolate assembled contigs. This pipeline takes assembled contigs as input and employs Prodigal [25] to predict genes from the contigs. These predicted genes are then aligned to the experimentally confirmed 485 MRGs in BacMet EXP MRG DB using DIAMOND with the alignment parameters: sequence identity ≥ 20%, value ≤ 1e-7, and alignment coverage ≥ 60% as discussed earlier in the “feature extraction” section. Then, using our feature computation approach, each predicted gene is represented by a vector *V* ∈ ℝ^66^ referring to the bit score-based similarity distribution with 66 types of BacMet EXP MRGs. Finally, DeepMRG is employed to identify and annotate MRGs. The entire pipeline was built using Nextflow [29], with parallel computation for the gene prediction step by Prodigal.

**Fig 6.**
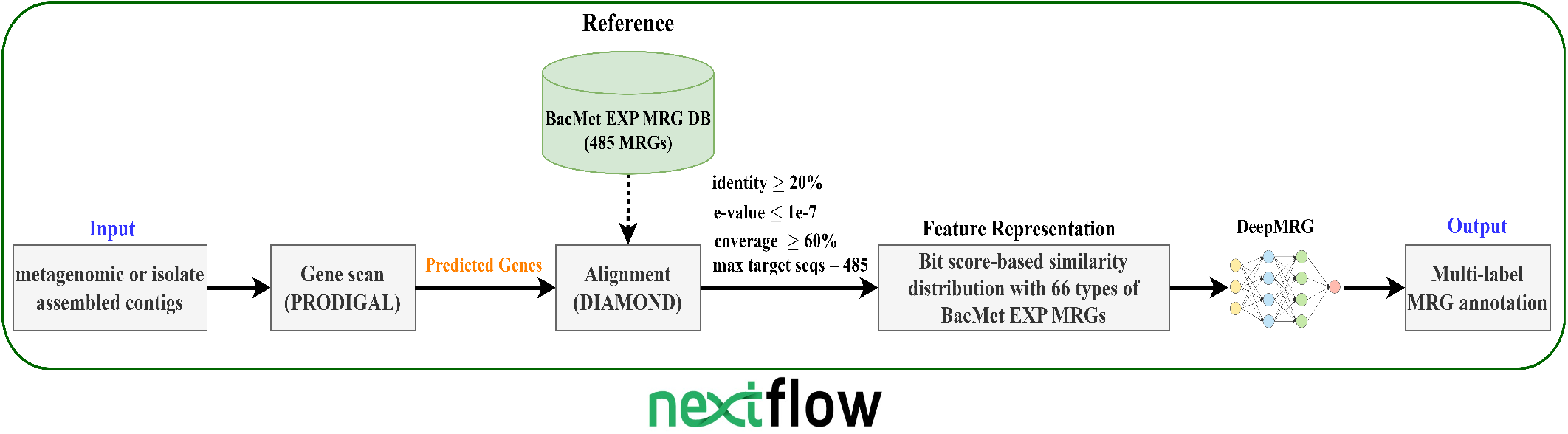
Pipeline for predicting bacterial MRGs from assembled contigs using DeepMRG.

## Discussion

In this paper, we developed a deep learning model, DeepMRG, for bacterial MRG classification. To the best of our knowledge, DeepMRG is the first work solely designed for bacterial MRG prediction which can offer detailed annotation for multi-metal resistance genes, indicating the specific metal labels to which the gene confers resistance. It leverages bit-score based similarity distribution with 66 types of experimentally confirmed MRGs. To mitigate the impact of sequence homology on model evaluation, we utilized a clustered split method with 40% identity threshold to create the training, validation, and test datasets. DeepMRG’s performance on the validation and test datasets demonstrates its ability to generalize well. DeepMRG performed better than AMRFinderPlus and the BLAST best hit methods on the test dataset. It outperformed AMRFinderPlus in identifying and classifying sequences in the BacMet Predicted MRG DB partition 2 dataset, where the sequences exhibit lower similarity (*<* 50%) to experimentally validated MRGs. Moreover, DeepMRG achieved comparable performance with the BLAST best hit method in this dataset. In the independent set of heavy metal resistance (HMR) genes from the *Cupriavidus* strain STM 6070, DeepMRG achieved better performance than AMRFinderPlus and the BLAST best hit method. Based on wet lab experiments, the strain STM 6070 exhibits heightened tolerance to Ni^2+^and Zn^2+^ concentrations. Our model, DeepMRG, demonstrated a high recall (≥ 0.75) in predicting the HMR genes of STM 6070 that confer resistance to these two metals. This validation underscores the effectiveness of our model in real-world scenarios. With the increasing prevalence of antimicrobial resistance worldwide, our tool can assist researchers in precisely and accurately identifying MRGs for effective mitigation of resistance spread.

Our model focuses entirely on MRG prediction. Users interested in multiple types of antimicrobial resistance (AMR), such as antibiotic drug and metal resistance, can combine our tool with others to obtain a more comprehensive AMR profile. For instance, if a gene sequence confers resistance to beta-lactams, copper, and zinc, users can employ DeepARG [17], HMD-ARG [30], ARG-SHINE [31], etc., for beta-lactam resistance prediction. DeepMRG can be utilized to predict metal resistances of the gene sequence.

## Availability and future directions

DeepMRG can be accessed through our web server at https://deepmrg.cs.vt.edu/deepmrg, providing a user-friendly interface for bacterial MRG prediction tasks. Alternatively, it can be installed locally and run as a command-line tool. The source code of DeepMRG, along with detailed instructions for installation and execution, can be found on our GitHub repository at https://github.com/muhit-emon/DeepMRG. All the datasets used in this paper are available from Zenodo (https://doi.org/10.5281/zenodo.10070602).

DeepMRG can process either full-length gene sequences or pre-assembled contigs for MRG prediction. In the case of contigs, the first step involves predicting open reading frames (ORFs), and subsequently, the resulting protein sequences are fed into DeepMRG for MRG prediction. However, the current pipeline of DeepMRG has a limitation in that it cannot directly handle short reads without the need for assembly. To address this limitation, our future plans involve expanding DeepMRG’s capabilities to enable direct prediction from short reads, eliminating the need for assembly. This development would significantly enhance DeepMRG’s applicability in genomics or metagenomics research and enable more efficient analysis of short read data.

As DeepMRG’s initial step involves aligning query gene sequences with experimentally validated MRGs, it might face some drawbacks typically associated with alignment-based methods. For example, DIAMOND may fail to identify MRGs very diverged from the experimentally validated MRGs, leading to false negatives. Comparatively, protein structures are more conserved than sequences and, therefore, can be leveraged to improve the performance of the prediction models. Consequently, we aim to incorporate protein 3D structures and protein language model (PLM) generated embeddings into DeepMRG. This integration of structural data and PLM is expected to significantly enhance the accuracy and reliability of the prediction model, providing deeper insights into the functional properties and mechanisms underlying metal resistance. By leveraging such information, DeepMRG will be better equipped to make precise predictions and contribute to a more comprehensive understanding of MRGs.

## Supporting information

**S1 File. DeepMRG documentation**. Complete documentation for using DeepMRG.

(PDF)

**S2 File. Types of the sequences in BacMet Predicted MRG DB partition 1**. Types of the sequences belonging to BacMet Predicted MRG DB partition 1 and the number of sequences within each type.

(PDF)

**S3 File. Types of BacMet EXP MRG DB sequences**. Types of the sequences in BacMet EXP MRG DB and the number of sequences within each type.

(PDF)

**S4 File. Performance evaluation metrics**. Details of the performance evaluation metrics used in our paper.

(PDF)

**S5 File. Detailed results of DeepMRG under 5-fold cross-validation**. Individual metal category-wise classification reports of DeepMRG for each fold under the 5-fold cross-validation experiment.

(PDF)

**S6 File. Detailed results of DeepMRG, AMRFinderPlus, and BLAST best hit approaches on BacMet Predicted MRG DB partition 2**. Individual metal category-wise classification reports of DeepMRG, AMRFinderPlus, and BLAST best hit methods (under different parameters) on the sequences in BacMet Predicted MRG DB partition 2.

(PDF)

**S7 File. BLAST best hit sequence identity of *Cupriavidus* strain STM 6070 heavy metal resistance (HMR) genes against the experimentally confirmed MRGs**.

BLAST best hits of all 53 STM 6070 HMR genes against the BacMet experimentally confirmed MRG database, including sequence identity and alignment coverage information.

(PDF)

**S8 File. Detailed results of DeepMRG, AMRFinderPlus with 50% identity threshold (AMRFinderPlus-50%), and BLAST best hit at 50% sequence identity cutoff (BLAST-50%) on the independent set**. (Table 1) Individual metal category-wise classification reports of DeepMRG, AMRFinderPlus-50%, and BLAST-50% on the HMR genes in the independent dataset. (Table 2) The overall F1-scores of DeepMRG (with the initial DIAMOND alignment parameters: identity ≥ 20%, e-value ≤ 1e-3, and alignment coverage ≥ 40%) on the independent dataset. (Table 3) Individual metal category-wise classification results of DeepMRG (with the initial DIAMOND alignment parameters: identity ≥ 20%, e-value ≤ 1e-3, and coverage ≥ 40%) on the independent set.

(PDF)

**S9 File. Gene Ontology (GO) terms of bacterial housekeeping genes**. Biological functions and GO terms associated with the bacterial housekeeping genes used to construct the negative microbial dataset.

(PDF)

**S1 Fig. Sequence identity histogram of BacMet Predicted MRG DB partition 2 against our training dataset**. Histogram of the best hit identity of the sequences in BacMet Predicted MRG DB partition 2 with our training dataset.

(TIF)

## Acknowledgments

We thank Professor Xiaofang Li from the Institute of Genetics and Developmental Biology of the Chinese Academy of Sciences for insightful suggestions.

## Author contributions

**Conceptualization:** Muhit Islam Emon, Liqing Zhang.

**Data curation:** Muhit Islam Emon.

**Formal analysis:** Muhit Islam Emon.

**Funding acquisition:** Liqing Zhang.

**Methodology:** Muhit Islam Emon.

**Project administration:** Liqing Zhang.

**Software & web server:** Muhit Islam Emon.

**Supervision:** Liqing Zhang.

**Validation:** Muhit Islam Emon.

**Writing – original draft:** Muhit Islam Emon.

**Writing – review & editing:** Muhit Islam Emon, Liqing Zhang.

