## Supplementary figures and images for "DeepMRG: a multi-label deep learning classifier for predicting bacterial metal resistance genes"

### S1 Fig

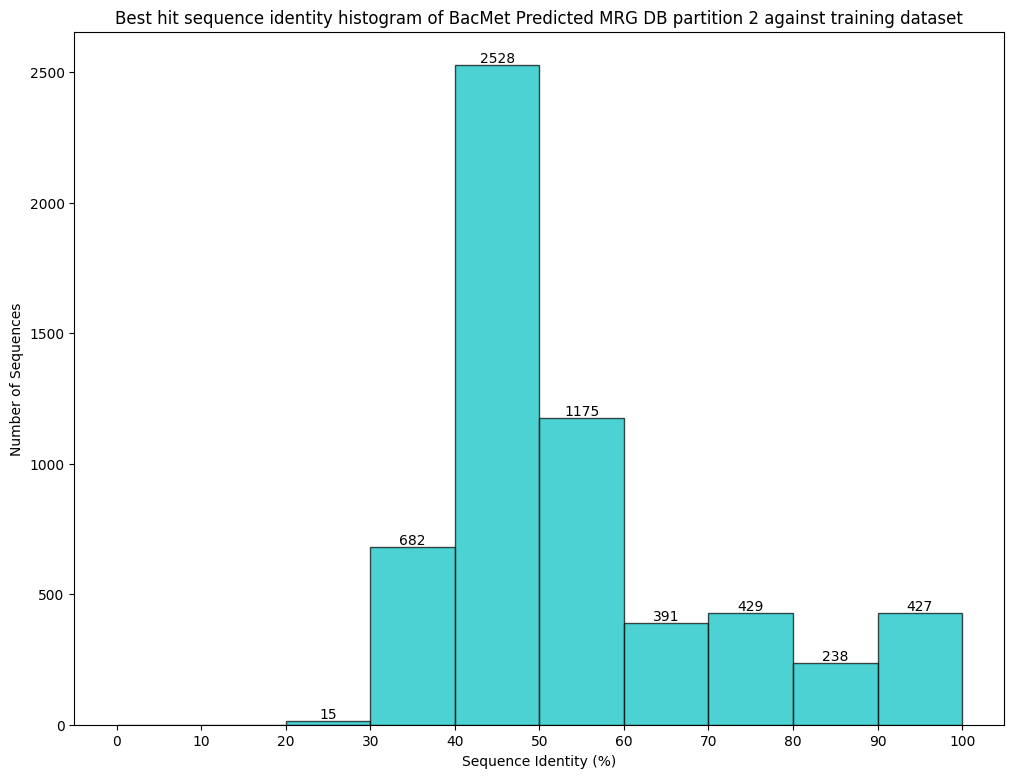
