## Supplementary material for "DeepMRG: a multi-label deep learning classifier for predicting bacterial metal resistance genes": S1 File

### S1 DeepMRG documentation

The web server of DeepMRG can be accessed at https://deepmrg.cs.vt.edu/deepmrg. DeepMRG can also be installed locally and used as a command line tool. The source code of DeepMRG is available at https://github.com/muhit-emon/DeepMRG. Here, we outline the instructions for using DeepMRG as a command line tool and via a web server. We also demonstrate how a user can test DeepMRG on supplied test datasets.

### 1. Using DeepMRG as a command line tool

#### Requirements

1. Linux operating system
2. conda

If conda is not installed on your machine, install conda for linux. To check if conda is installed successfully and can be accessed as a command, run the following:

**conda --version**

It will show the version of conda installed on your machine. Any version of conda works for using DeepMRG. The following figure shows an example of conda version checking.

```
(base) muhit@muhit-HP-Laptop-15-ef1xxx:~/Desktop$ conda --version
conda 23.7.2
```

#### Installation

To install DeepMRG do:

```
git clone https://github.com/muhit-emon/DeepMRG.git
cd DeepMRG
bash install.sh
```

Installation will take ~10 minutes as it creates a new conda environment and downloads binaries of diamond and prodigal softwares which are required to run DeepMRG.

#### Activation of conda environment

After installation of DeepMRG, a conda environment named **deepmrg** will be created. To activate the environment, run the following command:

```
conda activate deepmrg
```

The conda environment deepmrg contains all the packages that are required to run DeepMRG.

#### Classifying protein sequences in a fasta file

The **protein_pipeline.nf** script can be used to predict MRGs from protein sequences in a fasta file. To run DeepMRG on protein sequences to predict MRGs, go inside the DeepMRG directory and run the following command:

```
nextflow run protein_pipeline.nf --prot <absolute/path/to/protein/fasta/file>
--out_prefix <prefix of output file name>

rm -r work
```

The command line options for this script (**protein_pipeline.nf**) are:

- --`prot`: The absolute path of the fasta file containing protein sequences to be classified
- --`out_prefix`: The prefix of the output file name

An output tsv file named **<prefix of output file name>_DeepMRG_annotation.tsv** that contains MRG predictions by DeepMRG will be generated inside DeepMRG directory.

A demo usage to classify protein sequences in a fasta file is shown below:

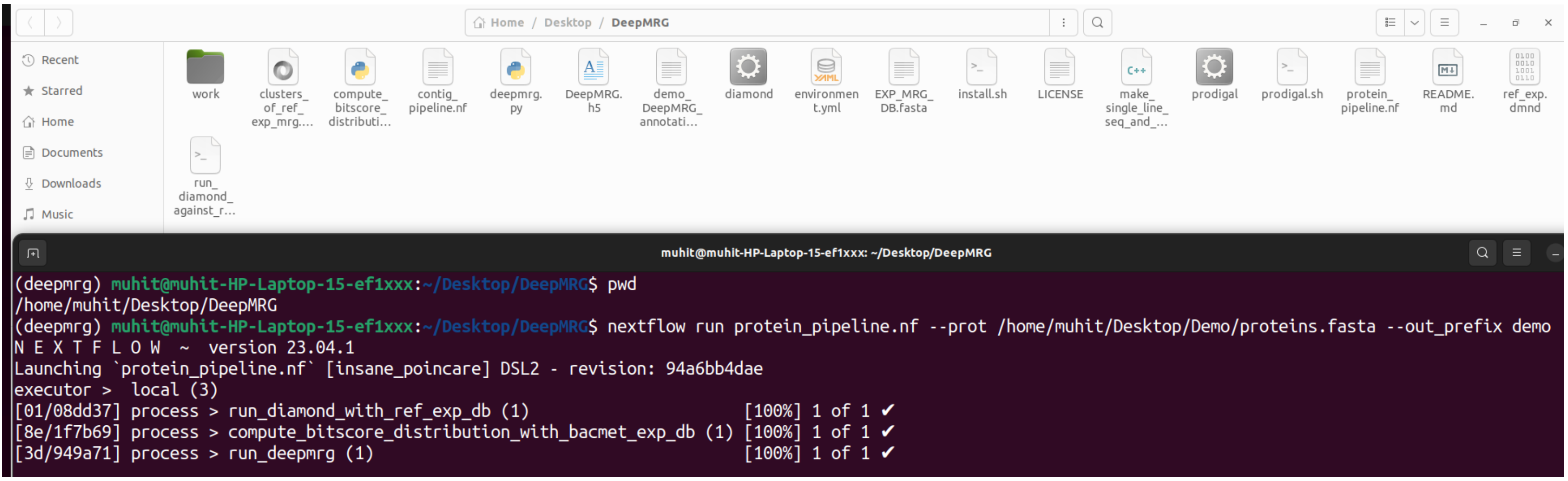


We executed the following command to use **protein_pipeline.nf** to classify the protein sequences in a fasta file:

```
nextflow run protein_pipeline.nf --prot /home/muhit/Desktop/Demo/proteins.fasta --out_prefix demo
```

Here, **/home/muhit/Desktop/Demo/proteins.fasta** is the absolute path of the fasta file that contains some protein sequences. We ran DeepMRG on this fasta file to predict MRGs. We used the string “demo” as --out_prefix.

The fasta file contains five protein sequences and it looks like the following:

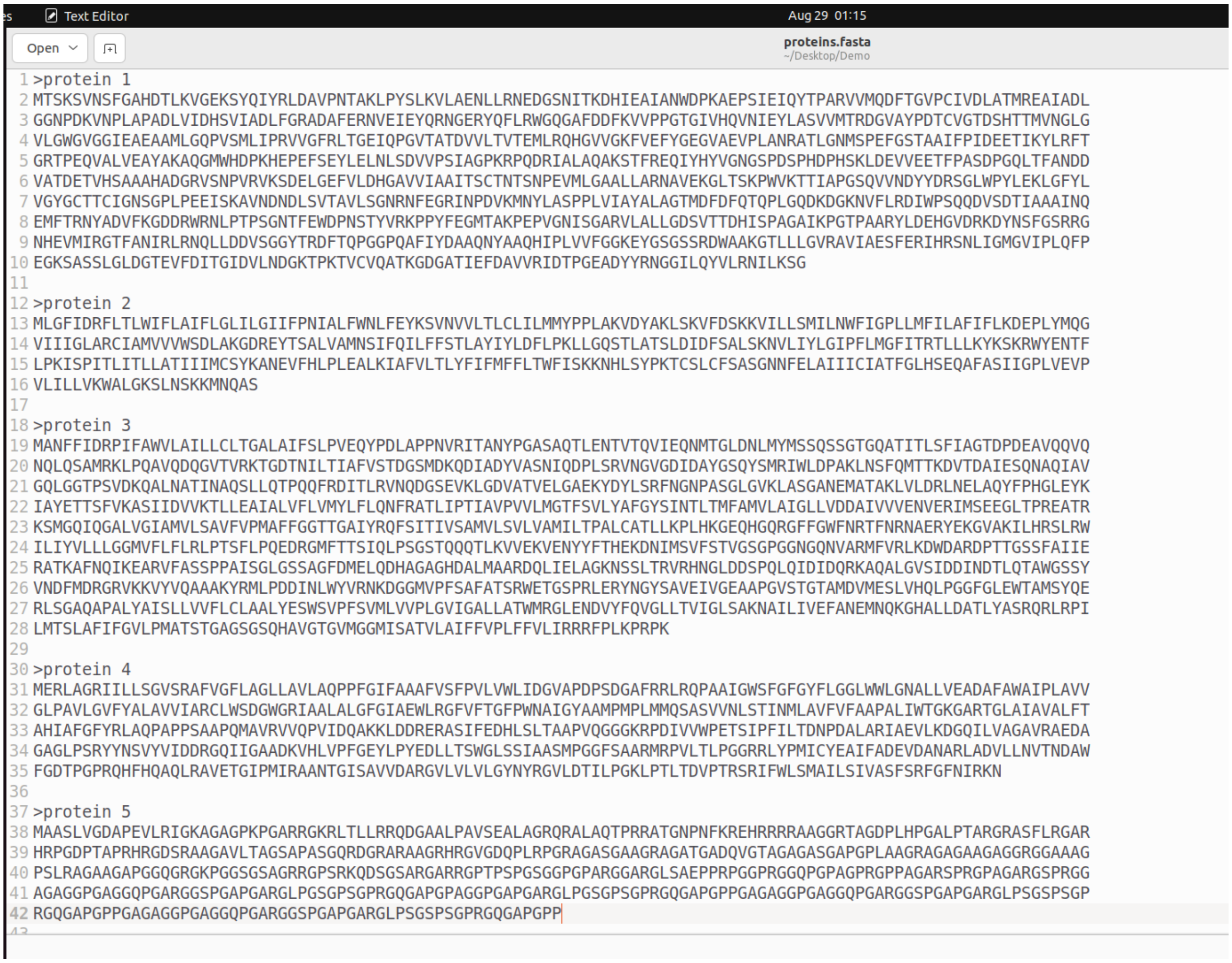


```
>protein 1
MTSKSVNSFGAHDTLKVGEKSYQIYRLDAVPNTAKLPYSLKVLAENLLRNEDGSNITKDHIEAIANWDPKAEPSIEIQYTPARVVMQDFTGVPCIVDLATMREAIADL
GGNPDKVNPLAPADLVIDHSVIADLFGRADAFERNVEIEYQRNGERYQFLRWGQGAFDDFKVVPPGTGIVHQVNIEYLASVVMTRDGVAYPDTCVGTDSHTTMVNGLG
VLGWGVGGIEAEAAMLGQPVSMLIPRVVGFRLTGEIQPGVTATDVVLTVTEMLRQHGVVGKFVEFYGEGVAEVPLANRATLGNMSPEFGSTAAIFPIDEETIKYLRFT
GRTPEQVALVEAYAKAQGMWHDPKHEPEFSEYLELNLSDVVPSIAGPKRPQDRIALAQAKSTFREQIYHYVGNGSPDSPHDPHSKLDEVVEETFPASDPGQLTFANDD
VATDETVHSAAAHADGRVSNPVRVKSDELGEFVLDHGAVVIAAITSCTNTSNPEVMLGAALLARNAVEKGLTSKPWVKTTIAPGSQVVNDYYDRSGLWPYLEKLGFYL
VGYGCTTCIGNSGPLPEEISKAVNDNDLSVTAVLSGNRNFEGRINPDVKMNYLASPPLVIAYALAGTMDFDFQTQPLGQDKDGKNVFLRDIWPSQQDVSDTIAAAINQ
EMFTRNYADVFKGDDRWRNLPTPSGNTFEWDPNSTYVRKPPYFEGMTAKPEPVGNISGARVLALLGDSVTTDHISPAGAIKPGTPAARYLDEHGVDRKDYNSFGSRRG
NHEVMIRGTFANIRLRNQLLDDVSGGYTRDFTQPGGPQAFIYDAAQNYAAQHIPLVVFGGKEYGSGSSRDWAAKGTLLLGVRAVIAESFERIHRSNLIGMGVIPLQFP
EGKSASSLGLDGTEVFDITGIDVLNDGKTPKTVCVQATKGDGATIEFDAVVRIDTPGEADYYRNGGILQYVLRNILKSG

>protein 2
MLGFIDRFLTLWIFLAIFLGLILGIIFPNIALFWNLFEYKSVNVVLTLCLILMMYPPLAKVDYAKLSKVFDSKKVILLSMILNWFIGPLLMFILAFIFLKDEPLYMQG
VIIIGLARCIAMVVVWSDLAKGDREYTSALVAMNSIFQILFFSTLAYIYLDFLPKLLGQSTLATSLDIDFSALSKNVLIYLGIPFLMGFITRTLLLKYKSKRWYENTF
LPKISPITLITLLATIIIMCSYKANEVFHLPLEALKIAFVLTLYFIFMFFLTWFISKKNHLSYPKTCSLCFSASGNNFELAIIICIATFGLHSEQAFASIIGPLVEVP
VLILLVKWALGKSLNSKKMNQAS

>protein 3
MANFFIDRPIFAWVLAILLCLTGALAIFSLPVEQYPDLAPPNVRITANYPGASAQTLENTVTQVIEQNMTGLDNLMYMSSQSSGTGQATITLSFIAGTDPDEAVQQVQ
NQLQSAMRKLPQAVQDQGVTVRKTGDTNILTIAFVSTDGSMDKQDIADYVASNIQDPLSRVNGVGDIDAYGSQYSMRIWLDPAKLNSFQMTTKDVTDAIESQNAQIAV
GQLGGTPSVDKQALNATINAQSLLQTPQQFRDITLRVNQDGSEVKLGDVATVELGAEKYDYLSRFNGNPASGLGVKLASGANEMATAKLVLDRLNELAQYFPHGLEYK
IAYETTSFVKASIIDVVKTLLEAIALVFLVMYLFLQNFRATLIPTIAVPVVLMGTFSVLYAFGYSINTLTMFAMVLAIGLLVDDAIVVVENVERIMSEEGLTPREATR
KSMGQIQGALVGIAMVLSAVFVPMAFFGGTTGAIYRQFSITIVSAMVLSVLVAMILTPALCATLLKPLHKGEQHGQRGFFGWFNRTFNRNAERYEKGVAKILHRSLRW
ILIYVLLLGGMVFLFLRLPTSFLPQEDRGMFTTSIQLPSGSTQQQTLKVVEKVENYYFTHEKDNIMSVFSTVGSGPGGNGQNVARMFVRLKDWDARDPTTGSSFAIIE
RATKAFNQIKEARVFASSPPAISGLGSSAGFDMELQDHAGAGHDALMAARDQLIELAGKNSSLTRVRHNGLDDSPQLQIDIDQRKAQALGVSIDDINDTLQTAWGSSY
VNDFMDRGRVKKVYVQAAAKYRMLPDDINLWYVRNKDGGMVPFSAFATSRWETGSPRLERYNGYSAVEIVGEAAPGVSTGTAMDVMESLVHQLPGGFGLEWTAMSYQE
RLSGAQAPALYAISLLVVFLCLAALYESWSVPFSVMLVVPLGVIGALLATWMRGLENDVYFQVGLLTVIGLSAKNAILIVEFANEMNQKGHALLDATLYASRQRLRPI
LMTSLAFIFGVLPMATSTGAGSGSQHAVGTGVMGGMISATVLAIFFVPLFFVLIRRRFPLKPRPK

>protein 4
MERLAGRIILLSGVSRAFVGFLAGLLAVLAQPPFGIFAAAFVSFPVLVWLIDGVAPDPSDGAFRRLRQPAAIGWSFGFGYFLGGLWWLGNALLVEADAFAWAIPLAVV
GLPAVLGVFYALAVVIARCLWSDGWGRIAALALGFGIAEWLRGFVFTGFPWNAIGYAAMPMPLMMQSASVVNLSTINMLAVFVFAAPALIWTGKGARTGLAIAVALFT
AHIAFGFYRLAQPAPPSAAPQMAVRVVQPVIDQAKKLDDRERASIFEDHLSLTAAPVQGGGKRPDIVVWPETSIPFILTDNPDALARIAEVLKDGQILVAGAVRAEDA
GAGLPSRYYNSVYVIDDRGQIIGAADKVHLVPFGEYLPYEDLLTSWGLSSIAASMPGGFSAARMRPVLTLPGGRRLYPMICYEAIFADEVDANARLADVLLNVTNDAW
FGDTPGPRQHFHQAQLRAVETGIPMIRAANTGISAVVDARGVLVLVLGYNYRGVLDTILPGKLPTLTDVPTRSRIFWLSMAILSIVASFSRFGFNIRKN

>protein 5
MAASLVGDAPEVLRIGKAGAGPKPGARRGKRLTLLRRQDGAALPAVSEALAGRQRALAQTPRRATGNPNFKREHRRRRAAGGRTAGDPLHPGALPTARGRASFLRGAR
HRPGDPTAPRHRGDSRAAGAVLTAGSAPASGQRDGRARAAGRHRGVGDQPLRPGRAGASGAAGRAGATGADQVGTAGAGASGAPGPLAAGRAGAGAAGAGGRGGAAAG
PSLRAGAAGAPGGQGRGKPGGSGSAGRRGPSRKQDSGSARGARRGPTPSPGSGGPGPARGGARGLSAEPPRPGGPRGGQPGPAGPRGPPAGARSPRGPAGARGSPRGG
AGAGGPGAGGQPGARGGSPGAPGARGLPGSGPSGPRGQGAPGPAGGPGAPGARGLPGSGPSGPRGQGAPGPPGAGAGGPGAGGQPGARGGSPGAPGARGLPSGSPSGP
RGQGAPGPPGAGAGGPGAGGQPGARGGSPGAPGARGLPSGSPSGPRGQGAPGPP
```

With **--out_prefix demo**, an output tsv file named **demo_DeepMRG_annotation.tsv** (contains MRG predictions by DeepMRG) has been generated inside DeepMRG directory.

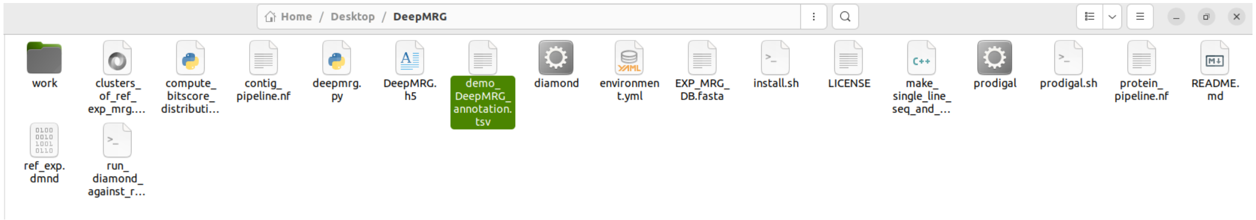


To remove the temporary work directory, do **rm -r work**

The output file is a tab separated file with each line containing a protein sequence header and the corresponding MRG predictions. The sequences are in the same order as in the input fasta file.

The output tsv (tab separated value) file **demo_DeepMRG_annotation.tsv** looks like the following:

| Protein_ID | Prediction |
|---|---|
| protein 1 | Iron (Fe) |
| protein 2 | Arsenic (As) |
| protein 3 | Copper (Cu),Zinc (Zn) |
| protein 4 | Copper (Cu),Zinc (Zn) |
| protein 5 | non-MRG |

The output file contains 2 columns:
The 1st column (**Protein_ID**) contains the header of the protein sequences in the input fasta file.
The 2nd column (**Prediction**) contains the prediction results of DeepMRG. By default, the proteins with the prediction score less than 3.5 (out of 5) are considered as non-MRG.

For example, protein 3 has been predicted to confer resistances to both Cu and Zn. On the other hand, protein 5 has been predicted as a non-MRG as its prediction score falls below 3.5.

#### Prediction of MRGs from metagenomic or isolate assembly (DNA sequences)

The **contig_pipeline.nf** script can be used to predict MRGs from metagenomic or isolate assembled contigs in a fasta file. To run DeepMRG on contigs to predict MRGs, go inside the DeepMRG directory and run the following command:

```
nextflow run contig_pipeline.nf --contig <absolute/path/to/contig/fasta/file>
--out_prefix <prefix of output file name>

rm -r work
```

The command line options for this script (**protein_pipeline.nf**) are:

- --`contig`: The absolute path of the fasta file containing contigs
- --`out_prefix`: The prefix of the output file name

The following two output files will be generated inside the DeepMRG directory:

1. **<prefix of output file name>_DeepMRG_annotation.tsv** (contains MRG predictions by DeepMRG).
2. **<prefix of output file name>_predicted_proteins.faa** (contains prodigal predicted proteins from contigs).

Given a contig file, we create multiple sub-files and run prodigal in parallel on these contig sub-files using nextflow to accelerate the gene prediction step. We output the prodigal predicted proteins from the contigs in the <prefix of output file name>_predicted_proteins.faa file.

A demo usage to run DeepMRG on contigs in a fasta file is shown below:

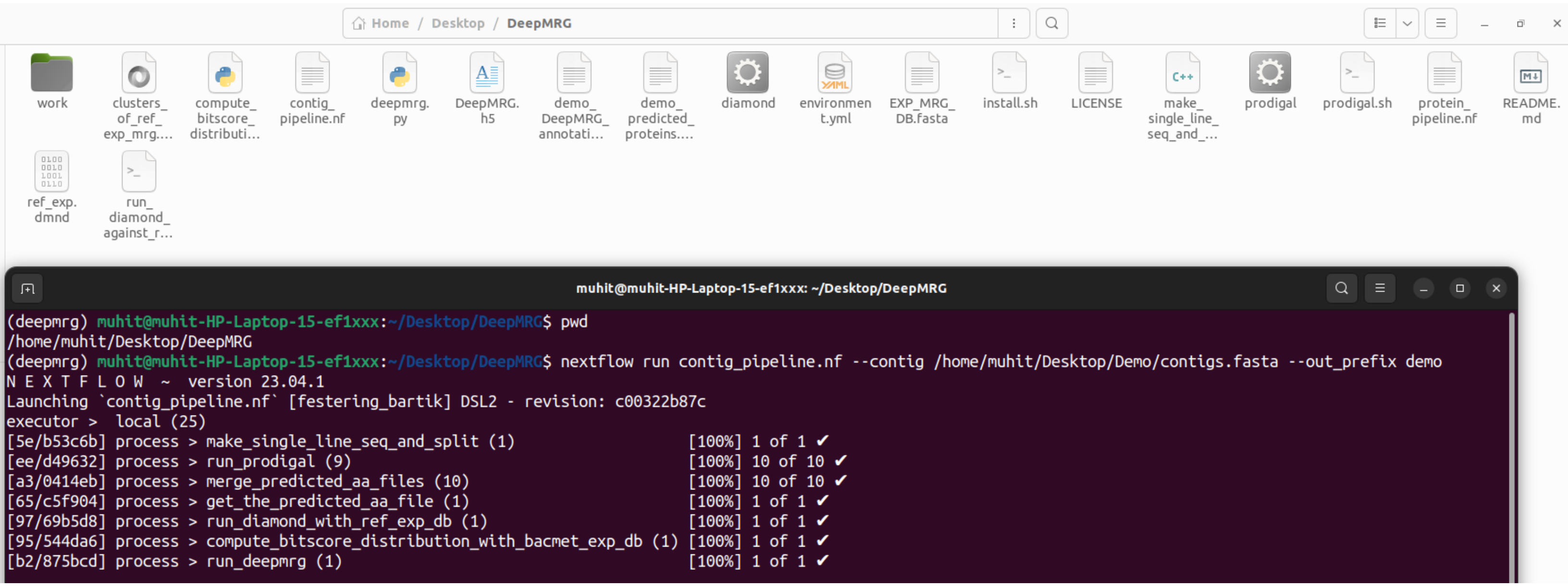


We executed the following command to use **contig_pipeline.nf** to predict MRGs from contigs in a fasta file:

nextflow run contig_pipeline.nf --contig /home/muhit/Desktop/Demo/contigs.fasta --out_prefix demo

Here, **/home/muhit/Desktop/Demo/contigs.fasta** is the absolute path of the fasta file that contains contigs (DNA sequences). We ran DeepMRG on this contig fasta file to predict MRGs. We used the string “demo” as --out_prefix.

A snippet of this contig fasta file is shown below:

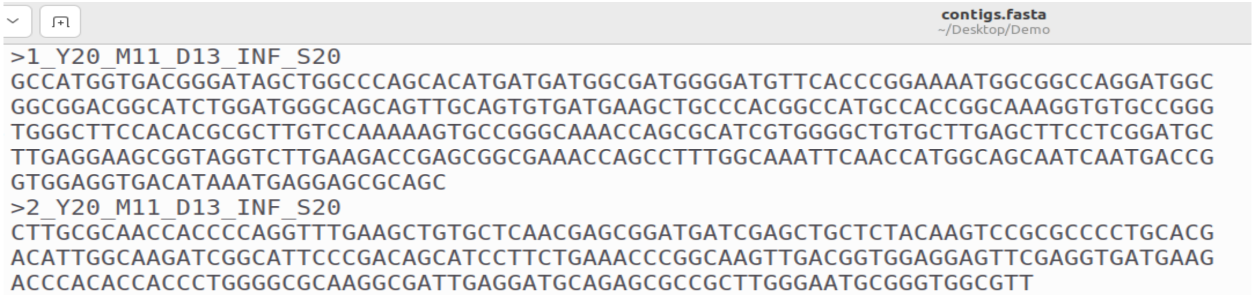

contigs.fasta
~/Desktop/Demo

```
>1_Y20_M11_D13_INF_S20
GCCATGGTGACGGGATAGCTGGCCCAGCACATGATGATGGCGATGGGGATGTTCACCCGGAAAATGGCGGCCAGGATGGC
GGCGGACGGCATCTGGATGGGCAGCAGTTGCAGTGTGATGAAGCTGCCCACGGCCATGCCACCGGCAAAGGTGTGCCGGG
TGGGCTTCCACACGCGCTTGTCCAAAAAGTGCCGGGCAAACCAGCGCATCGTGGGGCTGTGCTTGAGCTTCCTCGGATGC
TTGAGGAAGCGGTAGGTCTTGAAGACCGAGCGGCGAAACCAGCCTTTGGCAAATTCAACCATGGCAGCAATCAATGACCG
GTGGAGGTGACATAAATGAGGAGCGCAGC
>2_Y20_M11_D13_INF_S20
CTTGCGCAACCACCCCAGGTTTGAAGCTGTGCTCAACGAGCGGATGATCGAGCTGCTCTACAAGTCCGCGCCCCTGCACG
ACATTGGCAAGATCGGCATTCCCGACAGCATCCTTCTGAAACCCGGCAAGTTGACGGTGGAGGAGTTCGAGGTGATGAAG
ACCCACACCACCCTGGGGCGCAAGGCGATTGAGGATGCAGAGCGCCGCTTGGGAATGCGGGTGGCGTT
```

With **--out_prefix demo**, two output files have been generated inside the DeepMRG directory. One is **demo_DeepMRG_annotation.tsv** (contains MRG predictions by DeepMRG) and the other is **demo_predicted_proteins.faa** (contains prodigal predicted proteins from contigs).

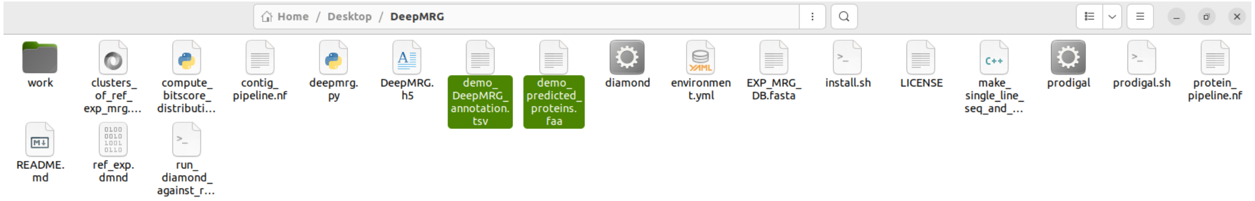


To remove the temporary work directory, do **rm -r work**.

A snippet of the prodigal predicted protein fasta file is shown below:

demo_predicted_proteins.faa
~/Desktop/DeepMRG

```
>501_Y20_M11_D13_INF_S20_1 # 3 # 599 # -1 # ID=1_1;partial=11;start_type=Edge;rbs_motif=None;rbs_spacer=None;gc_cont=0.434
IQNAALADRKSAWVNDGTVTIDPAVDKYLELAMQLIDNGYTNKNDPWTDGWQRDFTGDVF
GYFGCPWFVYWSINDEESGSATAGKRNICAGPSAYNWGGTYLSVTDKCPNKELAALVLKT
LCCDTDVMYKIQDETLDFVNNKAVVQKMIANGKGATPILGGANPLQTWYNVAVKVDGSKA
TQYDSVIDGYLYTVIDGCE
>502_Y20_M11_D13_INF_S20_1 # 3 # 104 # -1 # ID=2_1;partial=10;start_type=GTG;rbs_motif=None;rbs_spacer=None;gc_cont=0.696
MTGTVWFVGAGPGDPELITVKGRSLIERAGAILF
>502_Y20_M11_D13_INF_S20_2 # 611 # 2107 # 1 # ID=2_2;partial=01;start_type=ATG;rbs_motif=AGGA;rbs_spacer=5-10bp;gc_cont=0.731
MNRTAEDLLQELLSLGASRRIEAPGHARIDRAALRTLCAFANEPGQGGGYLLWGLEPVLP
DSAAPSAPGTGAAPTAPAVPSTWTAPATLADQAAAPASAVPQAQPAPDAPTTTPEPPPRY
RVVGVAQPEQLQAELARHCASAFNTALRPQVRTERLAGHTVVVAYLPEAAPATKPVFLAH
QGLPRGAYRRIGSSDQPGTEDDLAALYAARHATTLDAAVLPDAQWHDIDPAALEEYRQLR
RQAHPGAEELIWNDHDLLRALGCAQGEGEQLRPTVAGIVLFGTPLALRRCLPMARIDYIR
VPGRVWAPDPEAPYETQEIRAPLLAAIRRASNAVRYELTAPAPGTDSPAAHDDSLPQRAV
REAIVNAVLHRSYRLQAPVQIIRYDNRLEIRNPGHALKPSAQWDAPGALLRNPRIAAVLH
DVRLASTKGSGIRTLREQMLAHDLLPPTLASQRSADQFVATLLFHQFLGPDDRAWLRRLT
REALSDEEARALVLVRECG
>503_Y20_M11_D13_INF_S20_1 # 1 # 528 # 1 # ID=3_1;partial=10;start_type=Edge;rbs_motif=None;rbs_spacer=None;gc_cont=0.648
TIISVRRKTADGYDGAIGGDGQVTLGNIVVKGTARKVRKLYHGRVLAGFAGATADAFTLF
ERFEAKLEKHQGHLVRAAIELTRDWRTDRVLRRLEAMLAVADKDASLIITGNGDVLEPEN
GIVTIGSGGAYAQAAALALLNHSELPAVEIVRKSLEIAGELCIYTNMHHTIETLP*
```

The tsv output file **demo_DeepMRG_annotation.tsv** is a tab-separated file with each line containing a protein sequence header and the corresponding MRG predictions. The sequences are in the same order as in the prodigal predicted protein fasta file.

A snippet of **demo_DeepMRG_annotation.tsv** is shown below:

demo_DeepMRG_annotation.tsv
~/Desktop/DeepMRG

```
Protein_ID	Prediction
3001_Y20_M11_D13_INF_S20_1 # 62 # 394 # -1 # ID=1_1;partial=01;start_type=Edge;rbs_motif=None;rbs_spacer=None;gc_cont=0.318	non-MRG
3002_Y20_M11_D13_INF_S20_1 # 2 # 400 # -1 # ID=2_1;partial=10;start_type=GTG;rbs_motif=GGA/GAG/AGG;rbs_spacer=5-10bp;gc_cont=0.694	non-MRG
3002_Y20_M11_D13_INF_S20_2 # 403 # 876 # -1 # ID=2_2;partial=01;start_type=Edge;rbs_motif=None;rbs_spacer=None;gc_cont=0.700	non-MRG
3003_Y20_M11_D13_INF_S20_1 # 1 # 672 # -1 # ID=3_1;partial=11;start_type=Edge;rbs_motif=None;rbs_spacer=None;gc_cont=0.626	non-MRG
3004_Y20_M11_D13_INF_S20_1 # 2 # 328 # 1 # ID=4_1;partial=10;start_type=Edge;rbs_motif=None;rbs_spacer=None;gc_cont=0.618	non-MRG
3004_Y20_M11_D13_INF_S20_2 # 442 # 702 # 1 # ID=4_2;partial=01;start_type=ATG;rbs_motif=GGA/GAG/AGG;rbs_spacer=5-10bp;gc_cont=0.552	non-MRG
3006_Y20_M11_D13_INF_S20_1 # 1 # 375 # 1 # ID=6_1;partial=11;start_type=Edge;rbs_motif=None;rbs_spacer=None;gc_cont=0.704	non-MRG
3007_Y20_M11_D13_INF_S20_1 # 1 # 192 # 1 # ID=7_1;partial=10;start_type=Edge;rbs_motif=None;rbs_spacer=None;gc_cont=0.646	non-MRG
3007_Y20_M11_D13_INF_S20_2 # 222 # 1064 # 1 # ID=7_2;partial=01;start_type=ATG;rbs_motif=GGA/GAG/AGG;rbs_spacer=5-10bp;gc_cont=0.655	non-MRG
3008_Y20_M11_D13_INF_S20_1 # 1 # 309 # -1 # ID=8_1;partial=11;start_type=Edge;rbs_motif=None;rbs_spacer=None;gc_cont=0.440	non-MRG
3009_Y20_M11_D13_INF_S20_1 # 2 # 211 # -1 # ID=9_1;partial=10;start_type=ATG;rbs_motif=None;rbs_spacer=None;gc_cont=0.690	non-MRG
3009_Y20_M11_D13_INF_S20_2 # 211 # 321 # -1 # ID=9_2;partial=01;start_type=Edge;rbs_motif=None;rbs_spacer=None;gc_cont=0.622	non-MRG
3011_Y20_M11_D13_INF_S20_1 # 3 # 812 # 1 # ID=11_1;partial=10;start_type=Edge;rbs_motif=None;rbs_spacer=None;gc_cont=0.632	non-MRG
3012_Y20_M11_D13_INF_S20_1 # 5 # 226 # -1 # ID=12_1;partial=00;start_type=ATG;rbs_motif=GGA/GAG/AGG;rbs_spacer=5-10bp;gc_cont=0.437	non-MRG
3012_Y20_M11_D13_INF_S20_2 # 242 # 373 # -1 # ID=12_2;partial=01;start_type=Edge;rbs_motif=None;rbs_spacer=None;gc_cont=0.379	non-MRG
3013_Y20_M11_D13_INF_S20_1 # 182 # 739 # 1 # ID=13_1;partial=00;start_type=ATG;rbs_motif=AGGA;rbs_spacer=5-10bp;gc_cont=0.599	non-MRG
3013_Y20_M11_D13_INF_S20_2 # 754 # 864 # 1 # ID=13_2;partial=01;start_type=ATG;rbs_motif=AGGAG;rbs_spacer=5-10bp;gc_cont=0.586	non-MRG
3014_Y20_M11_D13_INF_S20_1 # 1 # 459 # 1 # ID=14_1;partial=11;start_type=Edge;rbs_motif=None;rbs_spacer=None;gc_cont=0.630	non-MRG
3015_Y20_M11_D13_INF_S20_1 # 2 # 775 # -1 # ID=15_1;partial=11;start_type=Edge;rbs_motif=None;rbs_spacer=None;gc_cont=0.745	non-MRG
3016_Y20_M11_D13_INF_S20_1 # 1 # 492 # 1 # ID=16_1;partial=11;start_type=Edge;rbs_motif=None;rbs_spacer=None;gc_cont=0.571	non-MRG
3017_Y20_M11_D13_INF_S20_1 # 1 # 378 # -1 # ID=17_1;partial=11;start_type=Edge;rbs_motif=None;rbs_spacer=None;gc_cont=0.664	non-MRG
3018_Y20_M11_D13_INF_S20_1 # 2 # 325 # -1 # ID=18_1;partial=11;start_type=Edge;rbs_motif=None;rbs_spacer=None;gc_cont=0.451	non-MRG
3019_Y20_M11_D13_INF_S20_1 # 2 # 289 # 1 # ID=19_1;partial=10;start_type=Edge;rbs_motif=None;rbs_spacer=None;gc_cont=0.552	non-MRG
3019_Y20_M11_D13_INF_S20_2 # 286 # 708 # 1 # ID=19_2;partial=01;start_type=GTG;rbs_motif=AGGAG;rbs_spacer=5-10bp;gc_cont=0.577	non-MRG
```

### 2. Using DeepMRG from the web server

The web server of DeepMRG can be accessed at https://deepmrg.cs.vt.edu/deepmrg.

#### Classifying protein sequences in a fasta file

**Step 1**: To predict MRGs from protein sequences in a fasta file, click on “Upload proteins” as shown below:

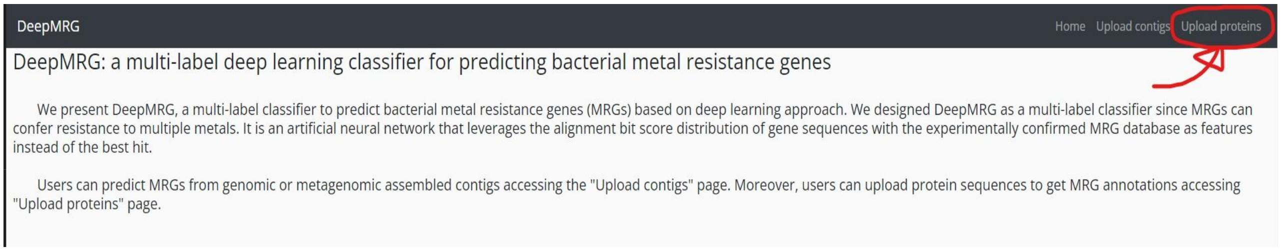


**Step 2**: Provide your email address (mandatory) and job name (mandatory) in the corresponding fields as shown below:

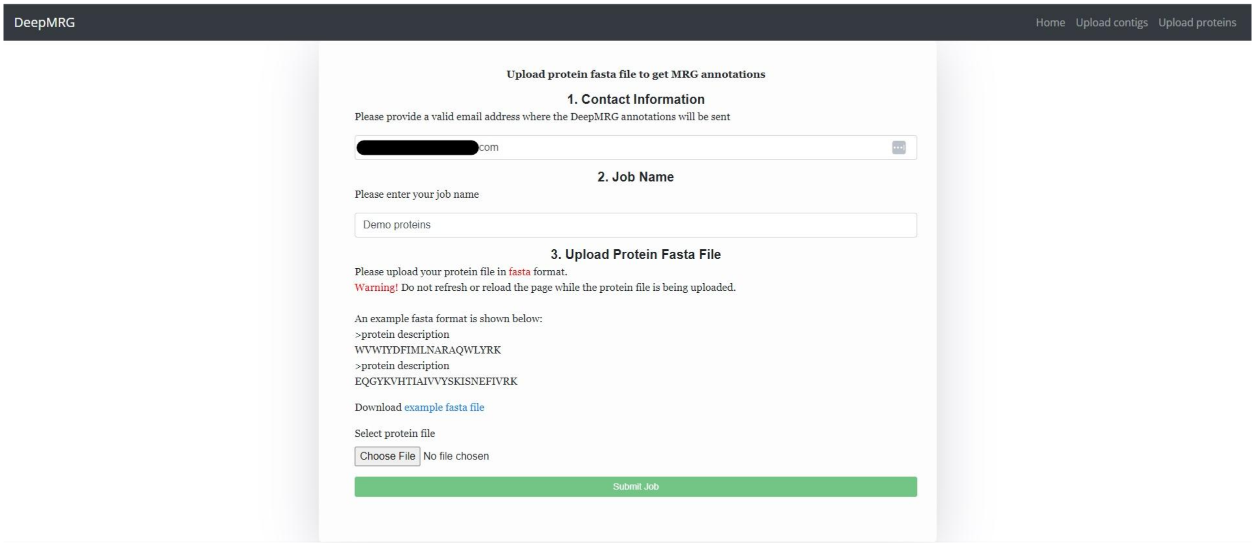


Here, we provided an email address (hidden) and “Demo proteins” as the job name.

**Step 3**: Select the **fasta file that contains the protein sequences** to be classified by DeepMRG. Here, we chose a fasta file named **proteins.fasta** to upload. The fasta file extension can be anything (*.fasta or *.fa or *.faa or *.txt). <u>Requirement is that the file should be in fasta format.</u>

After providing an email address, job name, and selecting the protein fasta file, we can submit the job to run DeepMRG.

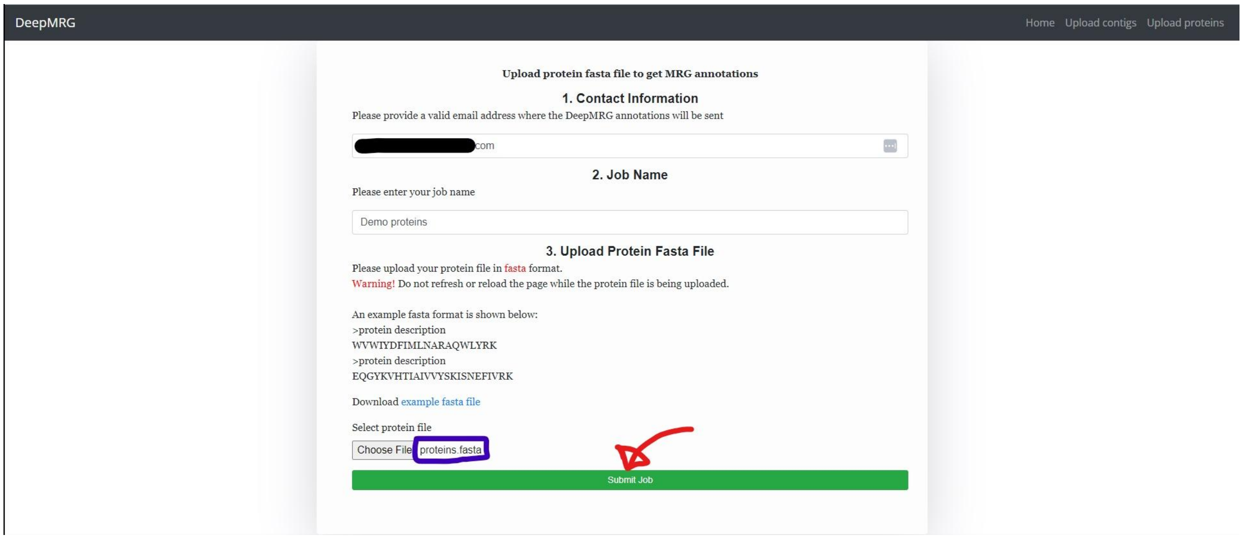


**Step 4**: If the file is uploaded successfully, we get a message as shown below:

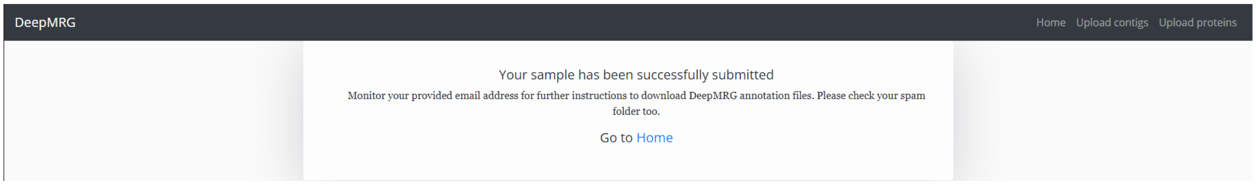


**Step 5**: When the job is done, we get an email in the provided email address as shown below:

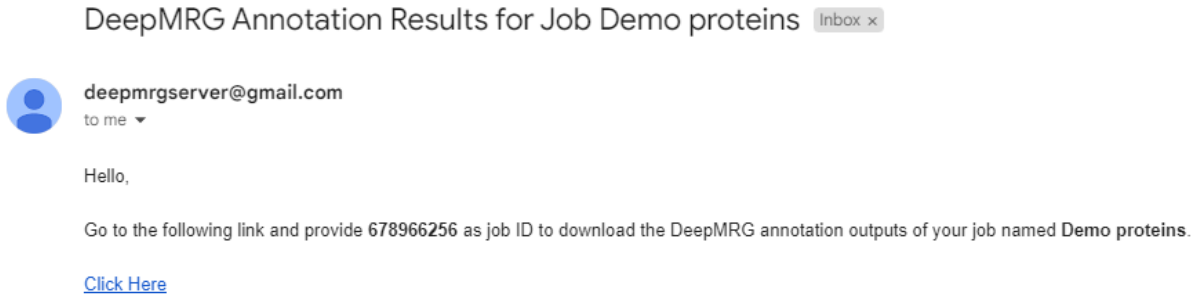


Clicking the “Click Here” link redirects to a web page. The provided job ID in the email is required to download the output file. For this example, the job ID is **678966256**.

After putting the job ID, we click the “Retrieve Download Link” button as shown below to create a link to download the output file. Then we click the link “Click here to download results” to download a zip file. The zip file name is **<job name>.zip**

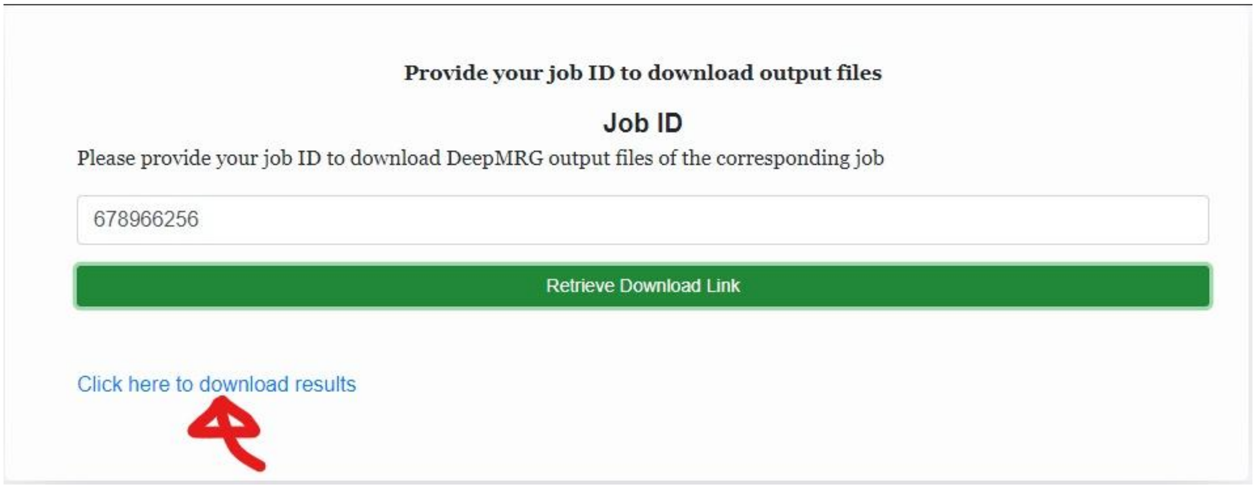


This compressed zip file contains a tsv file named **<job ID>_DeepMRG_annotation.tsv**

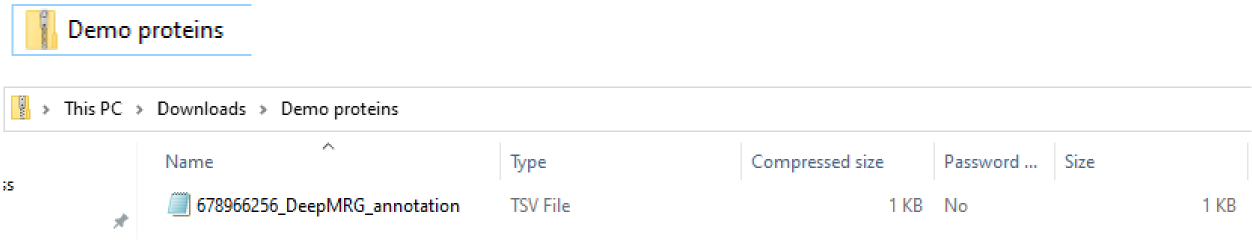


The output file **<job ID>_DeepMRG_annotation.tsv** is a tab separated file with each line containing a protein sequence header and the corresponding MRG predictions. The sequences are in the same order as in the input fasta file.

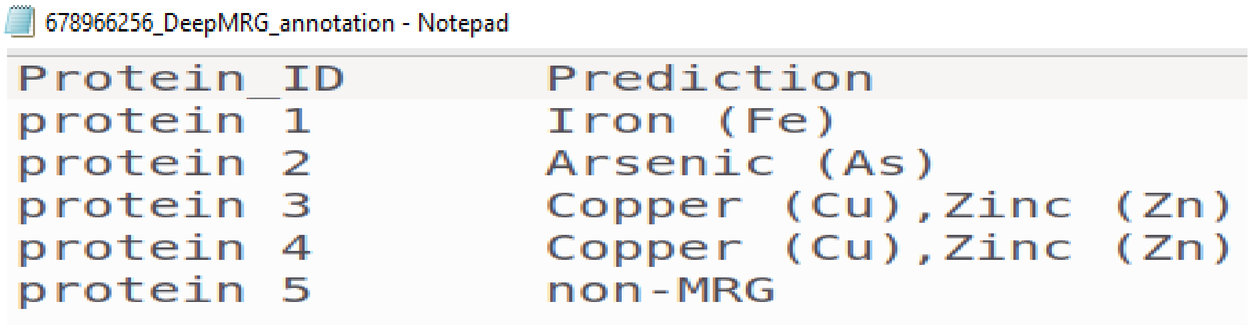
678966256_DeepMRG_annotation - Notepad

| Protein_ID | Prediction |
|---|---|
| protein 1 | Iron (Fe) |
| protein 2 | Arsenic (As) |
| protein 3 | Copper (Cu),Zinc (Zn) |
| protein 4 | Copper (Cu),Zinc (Zn) |
| protein 5 | non-MRG |

The output file contains 2 columns:
The 1st column (**Protein_ID**) contains the header of the protein sequences in the input fasta file.
The 2nd column (**Prediction**) contains the prediction results of DeepMRG. By default, the proteins with the prediction score less than 3.5 (out of 5) are regarded as non-MRG.

For example, protein 3 has been predicted to confer resistances to both Cu and Zn. On the other hand, protein 5 has been predicted as a non-MRG as its prediction score falls below 3.5.

#### Prediction of MRGs from metagenomic or isolate assembly (DNA sequences)

**Step 1**: To predict MRGs from (meta)genomic assembled contigs in a fasta file, click on "Upload contigs" as shown below:

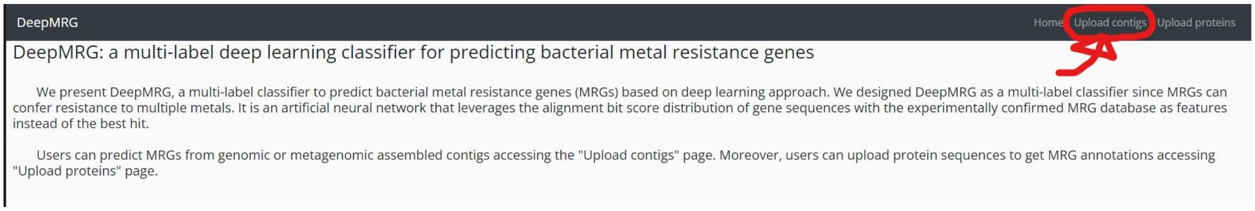


**Step 2**: Provide your email address (mandatory) and job name (mandatory) in the corresponding fields as shown below:

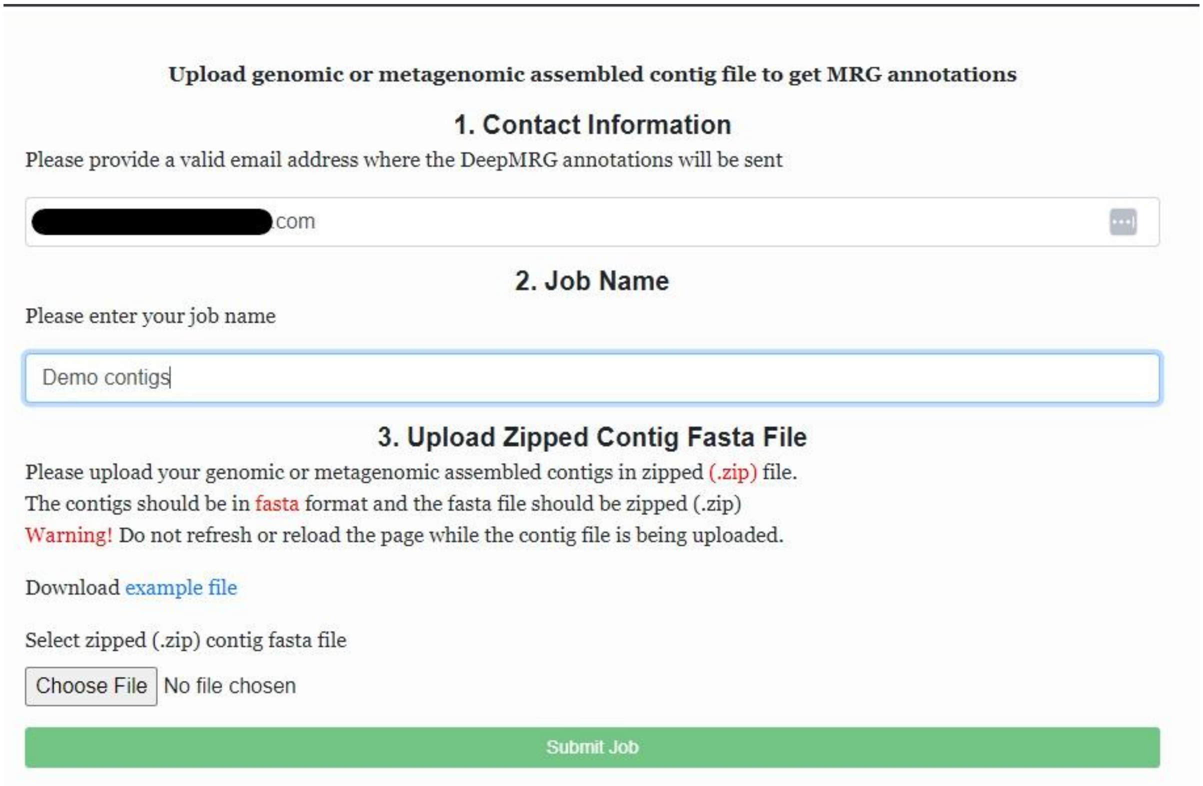

Here, we provided an email address (hidden) and “Demo contigs” as the job name.

**Step 3**: The genomic or metagenomic contigs must be in a fasta file. This fasta file is required to be zipped to upload to the web server. Therefore, if the fasta file containing the contigs is not zipped, zip it first and then submit the zipped fasta file to the web server.

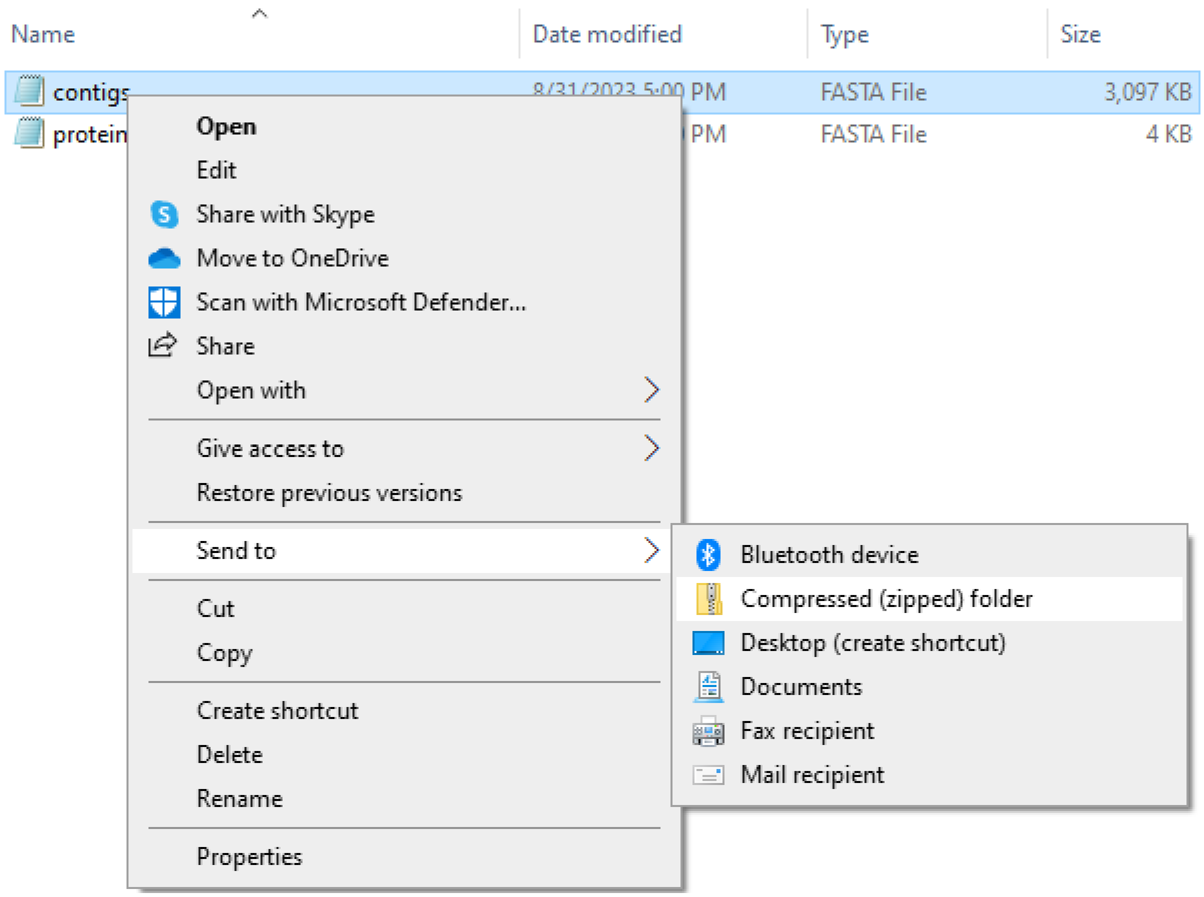


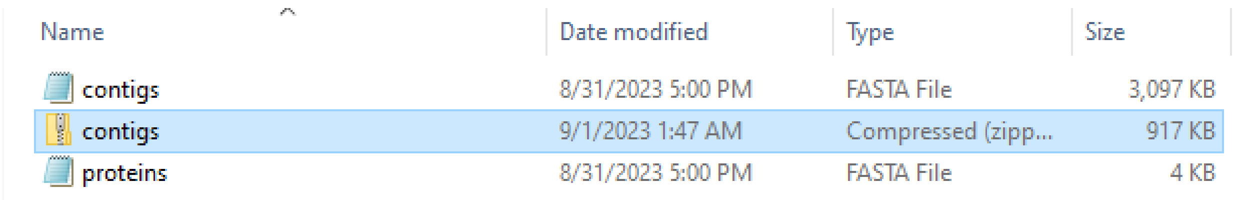


In Linux, we can compress a contig fasta file using the command **zip.** If your system doesn’t have zip installed, it can be installed by **sudo apt-get install zip.** We show an example of zipping contigs.fasta file on Ubuntu below:

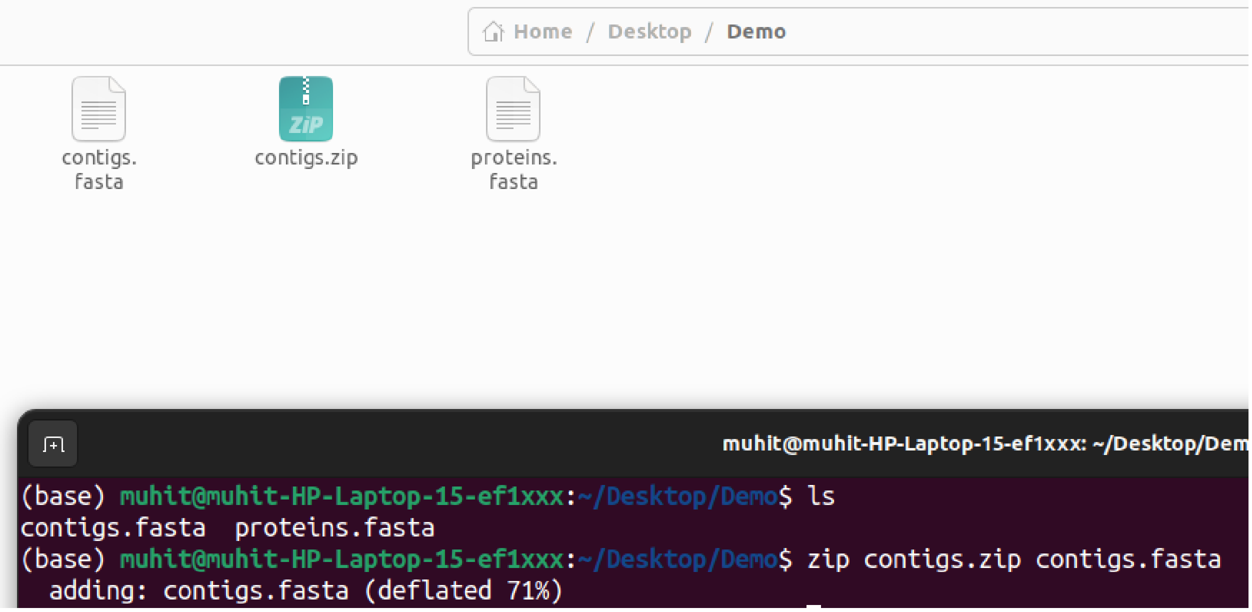


After providing an email address, job name, and selecting the zipped contig fasta file, we can submit the job to run DeepMRG.

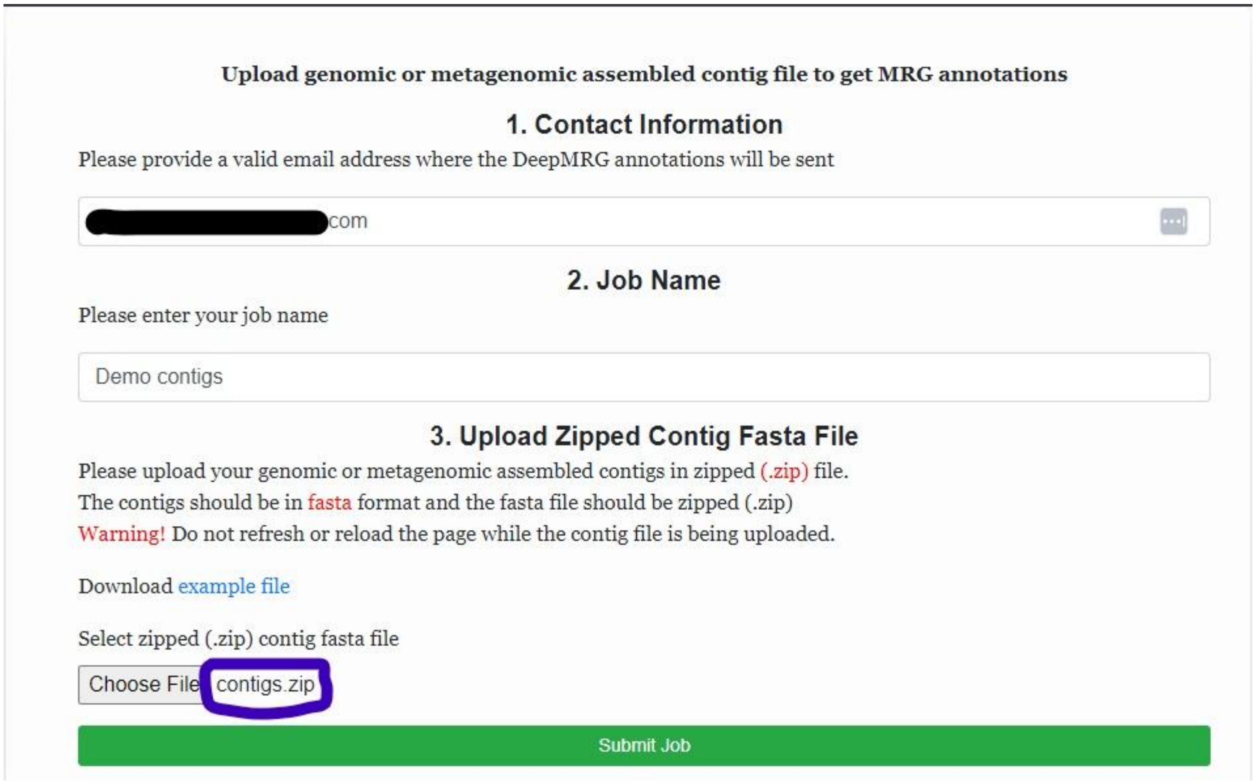

**Step 4**: If the file is uploaded successfully, we get a message as shown below:

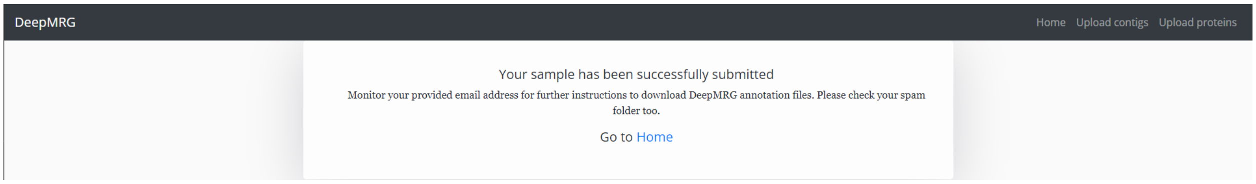


**Step 5**: When the job is done, we get an email in the provided email address as shown below:

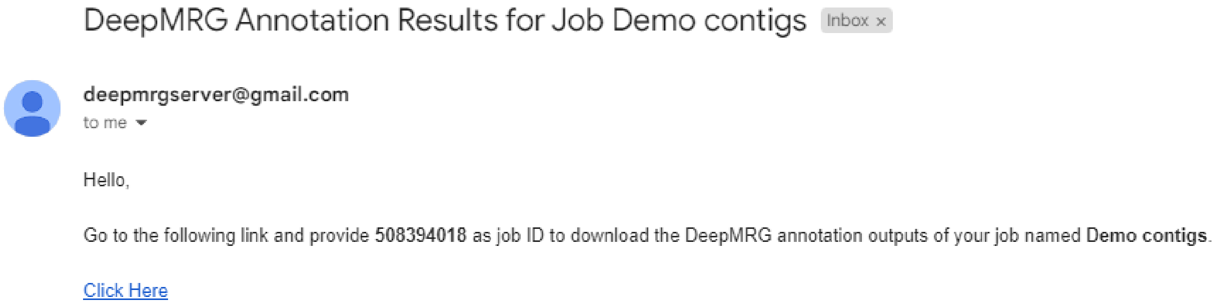


Clicking the “Click Here” link redirects to a web page. The provided job ID in the email is required to download the output file. For this example, the job ID is **508394018**.

After putting the job ID, we click the “Retrieve Download Link” button as shown below to create a link to download the output files. Then we click the link “Click here to download results” to download a zip file. The zip file name is **<job name>.zip**

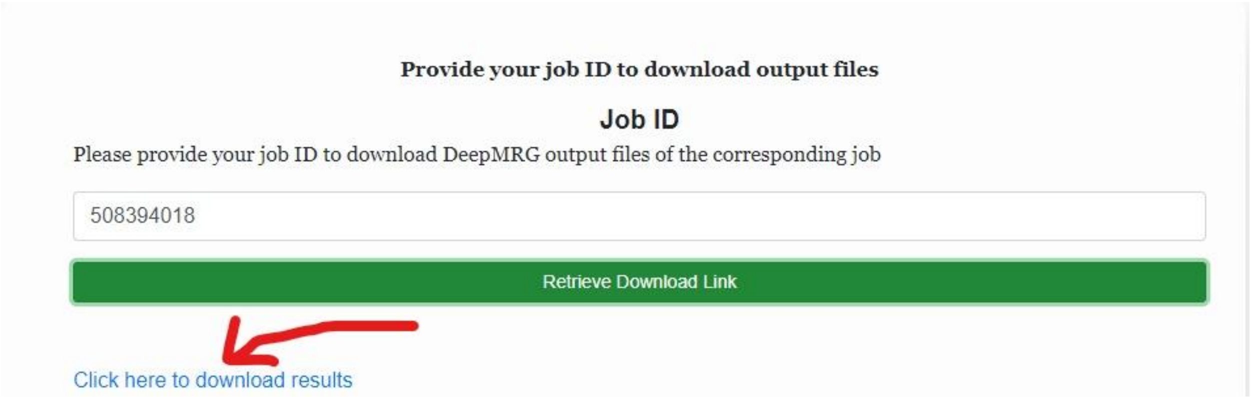

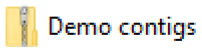


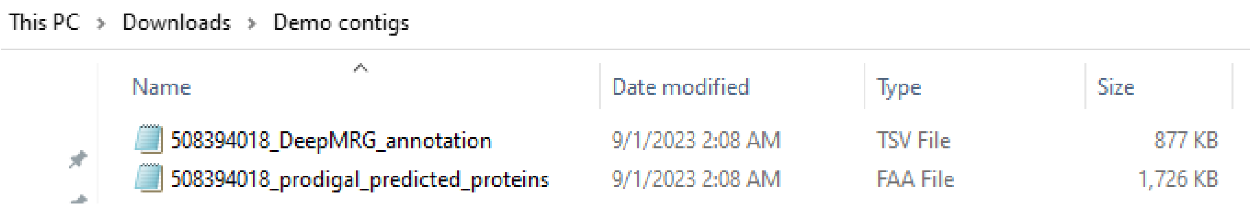


This compressed zip file contains two files:

1. **<job ID>_DeepMRG_annotation.tsv** (contains MRG predictions by DeepMRG)
2. **<job ID>_prodigal_predicted_proteins.faa** (contains prodigal predicted proteins from contigs)

The output file **<job ID>_DeepMRG_annotation.tsv** is a tab separated file with each line containing a protein sequence header and the corresponding MRG predictions. The sequences are in the same order as in the **<job ID>_prodigal_predicted_proteins.faa** fasta file.

### 3. Testing DeepMRG on supplied test datasets

The test datasets that we used to evaluate DeepMRG are available in our Github repository (https://github.com/muhit-emon/DeepMRG) under the **Test** directory. Additionally, we have provided Python and Nextflow scripts to assess DeepMRG and replicate the results using these test datasets.

We have provided the following test datasets in our Github repository:

- TEST.fasta
- BacMet_Predicted_MRG_DB_partition_2.fasta
- Fold1_Validation.fasta
- Fold2_Validation.fasta
- Fold3_Validation.fasta
- Fold4_Validation.fasta
- Fold5_Validation.fasta

At first, activate the conda environment and go inside the **Test** directory

```
conda activate deepmrg
cd DeepMRG/Test
```

#### Evaluation through 5-fold cross-validation

1. To evaluate DeepMRG on the 1st validation fold, execute the following command:

   ```
   nextflow run test_pipeline.nf --test_set fold1
   ```

   As output, a txt file named **fold1_classification_report.txt** will be generated inside the Test directory which contains the classification results of DeepMRG on **Fold1_Validation.fasta**

2. To evaluate DeepMRG on the 2nd validation fold, execute the following command:

   ```
   nextflow run test_pipeline.nf --test_set fold2
   ```

   As output, a txt file named **fold2_classification_report.txt** will be generated inside the Test directory which contains the classification results of DeepMRG on **Fold2_Validation.fasta**

3. To evaluate DeepMRG on the 3rd validation fold, execute the following command:

   ```
   nextflow run test_pipeline.nf --test_set fold3
   ```

   As output, a txt file named **fold3_classification_report.txt** will be generated inside the Test directory which contains the classification results of DeepMRG on **Fold3_Validation.fasta**

4. To evaluate DeepMRG on the 4th validation fold, execute the following command:

   ```
   nextflow run test_pipeline.nf --test_set fold4
   ```

   As output, a txt file named **fold4_classification_report.txt** will be generated inside the Test directory which contains the classification results of DeepMRG on **Fold4_Validation.fasta**

5. To evaluate DeepMRG on the 5th validation fold, execute the following command:

   ```
   nextflow run test_pipeline.nf --test_set fold5
   ```

As output, a txt file named **fold5_classification_report.txt** will be generated inside the Test directory which contains the classification results of DeepMRG on **Fold5_Validation.fasta**

#### Evaluation on test dataset (TEST.fasta)

Execute the following command:

```
nextflow run test_pipeline.nf --test_set TEST
```

As output, a txt file named **TEST_classification_report.txt** will be generated inside the Test directory which contains the classification results of DeepMRG on **TEST.fasta**

#### Evaluation on BacMet Predicted MRG DB partition 2

Execute the following command:

```
nextflow run test_pipeline.nf --test_set LOW
```

As output, a txt file named **LOW_classification_report.txt** will be generated inside the Test directory which contains the classification results of DeepMRG on **BacMet_Predicted_MRG_DB_partition_2.fasta**

#### Evaluation on the independent set of *Cupriavidus* strain STM 6070 HMR genes

Execute the following command:

```
nextflow run test_pipeline.nf --test_set IND
```

As output, a txt file named **IND_classification_report.txt** will be generated inside the Test directory which contains the classification results of DeepMRG on **STM6070_HMR_genes_IND.fasta**
