## Supplementary material for "DeepMRG: a multi-label deep learning classifier for predicting bacterial metal resistance genes": S2 File

### S2 Types of the sequences in BacMet Predicted MRG DB partition 1

| Type # | Type | Number of sequences |
|---|---|---|
| 1 | Aluminium (Al) | 718 |
| 2 | Antimony (Sb),Arsenic (As) | 1913 |
| 3 | Arsenic (As) | 5454 |
| 4 | Bismuth (Bi),Cadmium (Cd),Lead (Pb),Zinc (Zn) | 19 |
| 5 | Cadmium (Cd) | 1135 |
| 6 | Cadmium (Cd),Cobalt (Co),Copper (Cu),Gallium (Ga),Iron (Fe),Manganese (Mn),Nickel (Ni),Zinc (Zn) | 436 |
| 7 | Cadmium (Cd),Cobalt (Co),Copper (Cu),Iron (Fe),Nickel (Ni),Zinc (Zn) | 470 |
| 8 | Cadmium (Cd),Cobalt (Co),Iron (Fe),Manganese (Mn),Zinc (Zn) | 424 |
| 9 | Cadmium (Cd),Cobalt (Co),Iron (Fe),Nickel (Ni),Zinc (Zn) | 475 |
| 10 | Cadmium (Cd),Cobalt (Co),Nickel (Ni) | 15 |
| 11 | Cadmium (Cd),Cobalt (Co),Nickel (Ni),Zinc (Zn) | 8 |
| 12 | Cadmium (Cd),Cobalt (Co),Zinc (Zn) | 39 |
| 13 | Cadmium (Cd),Lead (Pb),Nickel (Ni) | 452 |
| 14 | Cadmium (Cd),Lead (Pb),Zinc (Zn) | 499 |
| 15 | Cadmium (Cd),Manganese (Mn) | 390 |
| 16 | Cadmium (Cd),Mercury (Hg) | 620 |
| 17 | Cadmium (Cd),Mercury (Hg),Silver (Ag) | 486 |
| 18 | Cadmium (Cd),Mercury (Hg),Zinc (Zn) | 427 |
| 19 | Cadmium (Cd),Nickel (Ni),Zinc (Zn) | 9 |
| 20 | Cadmium (Cd),Zinc (Zn) | 2599 |
| 21 | Chromium (Cr) | 1870 |
| 22 | Chromium (Cr),Iron (Fe) | 461 |
| 23 | Chromium (Cr),Molybdenum (Mo),Vanadium (V) | 444 |

| | | |
|---|---|---|
| 24 | Chromium (Cr),Selenium (Se),Tellurium (Te) | 1257 |
| 25 | Cobalt (Co) | 2 |
| 26 | Cobalt (Co),Copper (Cu) | 1006 |
| 27 | Cobalt (Co),Gallium (Ga),Iron (Fe),Nickel (Ni) | 456 |
| 28 | Cobalt (Co),Iron (Fe),Nickel (Ni) | 750 |
| 29 | Cobalt (Co),Magnesium (Mg) | 1781 |
| 30 | Cobalt (Co),Magnesium (Mg),Manganese (Mn),Nickel (Ni) | 428 |
| 31 | Cobalt (Co),Nickel (Ni) | 1955 |
| 32 | Cobalt (Co),Nickel (Ni),Zinc (Zn) | 1 |
| 33 | Copper (Cu) | 15783 |
| 34 | Copper (Cu),Gold (Au) | 458 |
| 35 | Copper (Cu),Iron (Fe),Manganese (Mn) | 235 |
| 36 | Copper (Cu),Nickel (Ni),Zinc (Zn) | 47 |
| 37 | Copper (Cu),Silver (Ag) | 3000 |
| 38 | Copper (Cu),Tellurium (Te) | 20 |
| 39 | Copper (Cu),Zinc (Zn) | 788 |
| 40 | Gallium (Ga),Iron (Fe) | 390 |
| 41 | Gold (Au) | 812 |
| 42 | Iron (Fe) | 5099 |
| 43 | Iron (Fe),Manganese (Mn) | 1734 |
| 44 | Iron (Fe),Manganese (Mn),Zinc (Zn) | 19 |
| 45 | Iron (Fe),Nickel (Ni) | 487 |
| 46 | Lead (Pb) | 21 |
| 47 | Lead (Pb),Zinc (Zn) | 441 |
| 48 | Magnesium (Mg),Manganese (Mn) | 802 |
| 49 | Manganese (Mn),Zinc (Zn) | 462 |

| | | |
|---|---|---|
| 50 | Mercury (Hg) | 2893 |
| 51 | Mercury (Hg),Zinc (Zn) | 164 |
| 52 | Molybdenum (Mo),Tungsten (W) | 3317 |
| 53 | Molybdenum (Mo),Tungsten (W),Vanadium (V) | 1 |
| 54 | Nickel (Ni) | 3249 |
| 55 | Nickel (Ni),Zinc (Zn) | 159 |
| 56 | Selenium (Se) | 909 |
| 57 | Silver (Ag) | 1542 |
| 58 | Tellurium (Te) | 1145 |
| 59 | Tellurium (Te),Zinc (Zn) | 468 |
| 60 | Tungsten (W) | 173 |
| 61 | Tungsten (W),Zinc (Zn) | 814 |
| 62 | Vanadium (V) | 105 |
| 63 | Zinc (Zn) | 5476 |
