## Supplementary material for "DeepMRG: a multi-label deep learning classifier for predicting bacterial metal resistance genes": S3 File

### S3 Types of BacMet EXP MRG DB sequences

| Type # | Type | Number of sequences |
|---|---|---|
| 1 | Aluminium (Al) | 2 |
| 2 | Antimony (Sb),Arsenic (As) | 16 |
| 3 | Antimony (Sb),Arsenic (As),Bismuth (Bi) | 4 |
| 4 | Arsenic (As) | 28 |
| 5 | Bismuth (Bi),Cadmium (Cd),Lead (Pb),Zinc (Zn) | 1 |
| 6 | Cadmium (Cd) | 7 |
| 7 | Cadmium (Cd),Cobalt (Co),Copper (Cu),Gallium (Ga),Iron (Fe),Manganese (Mn),Nickel (Ni),Zinc (Zn) | 1 |
| 8 | Cadmium (Cd),Cobalt (Co),Copper (Cu),Iron (Fe),Nickel (Ni),Zinc (Zn) | 1 |
| 9 | Cadmium (Cd),Cobalt (Co),Iron (Fe),Manganese (Mn),Zinc (Zn) | 1 |
| 10 | Cadmium (Cd),Cobalt (Co),Iron (Fe),Nickel (Ni),Zinc (Zn) | 1 |
| 11 | Cadmium (Cd),Cobalt (Co),Nickel (Ni) | 8 |
| 12 | Cadmium (Cd),Cobalt (Co),Nickel (Ni),Zinc (Zn) | 1 |
| 13 | Cadmium (Cd),Cobalt (Co),Zinc (Zn) | 8 |
| 14 | Cadmium (Cd),Lead (Pb),Nickel (Ni) | 1 |
| 15 | Cadmium (Cd),Lead (Pb),Zinc (Zn) | 1 |
| 16 | Cadmium (Cd),Manganese (Mn) | 1 |
| 17 | Cadmium (Cd),Mercury (Hg) | 2 |
| 18 | Cadmium (Cd),Mercury (Hg),Silver (Ag) | 1 |
| 19 | Cadmium (Cd),Mercury (Hg),Zinc (Zn) | 1 |
| 20 | Cadmium (Cd),Nickel (Ni),Zinc (Zn) | 3 |
| 21 | Cadmium (Cd),Zinc (Zn) | 11 |
| 22 | Chromium (Cr) | 18 |
| 23 | Chromium (Cr),Cobalt (Co),Zinc (Zn) | 1 |
| 24 | Chromium (Cr),Iron (Fe) | 1 |
| 25 | Chromium (Cr),Molybdenum (Mo),Vanadium (V) | 1 |
| 26 | Chromium (Cr),Selenium (Se),Tellurium (Te) | 2 |
| 27 | Cobalt (Co) | 2 |
| 28 | Cobalt (Co),Copper (Cu) | 4 |
| 29 | Cobalt (Co),Gallium (Ga),Iron (Fe),Nickel (Ni) | 1 |
| 30 | Cobalt (Co),Iron (Fe),Nickel (Ni) | 2 |

| | | |
|---|---|---|
| 31 | Cobalt (Co),Magnesium (Mg) | 6 |
| 32 | Cobalt (Co),Magnesium (Mg),Manganese (Mn),Nickel (Ni) | 2 |
| 33 | Cobalt (Co),Nickel (Ni) | 29 |
| 34 | Cobalt (Co),Nickel (Ni),Zinc (Zn) | 1 |
| 35 | Copper (Cu) | 109 |
| 36 | Copper (Cu),Gold (Au) | 1 |
| 37 | Copper (Cu),Iron (Fe),Manganese (Mn) | 1 |
| 38 | Copper (Cu),Nickel (Ni),Zinc (Zn) | 1 |
| 39 | Copper (Cu),Silver (Ag) | 10 |
| 40 | Copper (Cu),Tellurium (Te) | 1 |
| 41 | Copper (Cu),Tungsten (W),Zinc (Zn) | 2 |
| 42 | Copper (Cu),Zinc (Zn) | 2 |
| 43 | Gallium (Ga),Iron (Fe) | 3 |
| 44 | Gold (Au) | 4 |
| 45 | Iron (Fe) | 14 |
| 46 | Iron (Fe),Manganese (Mn) | 8 |
| 47 | Iron (Fe),Manganese (Mn),Zinc (Zn) | 5 |
| 48 | Iron (Fe),Nickel (Ni) | 2 |
| 49 | Lead (Pb) | 5 |
| 50 | Lead (Pb),Zinc (Zn) | 1 |
| 51 | Magnesium (Mg),Manganese (Mn) | 2 |
| 52 | Manganese (Mn),Zinc (Zn) | 1 |
| 53 | Mercury (Hg) | 59 |
| 54 | Mercury (Hg),Zinc (Zn) | 1 |
| 55 | Molybdenum (Mo),Tungsten (W) | 11 |
| 56 | Molybdenum (Mo),Tungsten (W),Vanadium (V) | 1 |
| 57 | Nickel (Ni) | 22 |
| 58 | Nickel (Ni),Zinc (Zn) | 1 |
| 59 | Selenium (Se) | 2 |
| 60 | Silver (Ag) | 8 |
| 61 | Tellurium (Te) | 13 |
| 62 | Tellurium (Te),Zinc (Zn) | 1 |
| 63 | Tungsten (W) | 3 |
| 64 | Tungsten (W),Zinc (Zn) | 2 |

| | | |
|---|---|---|
| 65 | Vanadium (V) | 1 |
| 66 | Zinc (Zn) | 18 |
