## Supplementary material for "DeepMRG: a multi-label deep learning classifier for predicting bacterial metal resistance genes": S4 File

### S4 Multi-label classification performance evaluation metrics

#### i. Label-based metrics

For the j[th] class, precision, recall, and F1-score are defined as follows:

$$Precision_j = \frac{TP_j}{TP_j+FP_j} \tag{1}$$

$$Recall_j = \frac{TP_j}{TP_j+FN_j} \tag{2}$$

$$F1_j = \frac{2\times TP_j}{2\times TP_j+FN_j+FP_j} \tag{3}$$

where $TP_j$, $FP_j$, and $FN_j$ represent the number of true positives, false positives, and false negatives respectively for the j[th] class.

Based on these quantities, macro-average precision, macro-average recall, and macro-average F1-score are defined as follows:

$$macro - avg\ Precision = \frac{1}{q}\sum_{i=1}^{q} Precision_j \tag{4}$$

$$macro - avg\ Recall = \frac{1}{q}\sum_{i=1}^{q} Recall_j \tag{5}$$

$$macro - avg\ F1 = \frac{1}{q}\sum_{i=1}^{q} F1_j \tag{6}$$

where $q$ is the number of class labels in the dataset.

#### ii. Sample-based metrics

$$samples - avg\ Precision = \frac{1}{p}\sum_{i=1}^{P} \frac{|Y_i \cap \hat{Y}_i|}{|\hat{Y}_i|} \tag{7}$$

$$samples - avg\ Recall = \frac{1}{p}\sum_{i=1}^{P} \frac{|Y_i \cap \hat{Y}_i|}{|Y_i|} \tag{8}$$

$$samples - avg\ F1 = \frac{1}{p}\sum_{i=1}^{P} \frac{2\times Precision_{sample}\times Recall_{sample}}{Precision_{sample}+Recall_{sample}} \tag{9}$$

where $p$ is the total number of samples in the dataset. For the i[th] sample, $Y_i$ denotes the ground-truth label set, while $\hat{Y}_i$ denotes the set of predicted labels.
