## Supplementary material for "DeepMRG: a multi-label deep learning classifier for predicting bacterial metal resistance genes": S5 File

### S5 Detailed results of DeepMRG under 5-fold cross-validation

Performance of DeepMRG for each individual metal label available in 1st fold

| Metal | Precision | Recall | F1-score | # of sequences |
|---|---|---|---|---|
| Arsenic (As) | 1 | 1 | 1 | 517 |
| Cadmium (Cd) | 1 | 1 | 1 | 839 |
| Chromium (Cr) | 1 | 1 | 1 | 62 |
| Cobalt (Co) | 1 | 1 | 1 | 7 |
| Copper (Cu) | 0.88 | 1 | 0.93 | 1582 |
| Iron (Fe) | 1 | 1 | 1 | 842 |
| Mercury (Hg) | 1 | 1 | 1 | 14 |
| Molybdenum (Mo) | 1 | 1 | 1 | 40 |
| Nickel (Ni) | 1 | 1 | 1 | 473 |
| Silver (Ag) | 1 | 1 | 1 | 1289 |
| Tellurium (Te) | 1 | 1 | 1 | 135 |
| Tungsten (W) | 1 | 1 | 1 | 40 |
| Zinc (Zn) | 1 | 1 | 1 | 2235 |

Performance of DeepMRG for each individual metal label available in 2nd fold

| Metal | Precision | Recall | F1-score | # of sequences |
|---|---|---|---|---|
| Antimony (Sb) | 1 | 1 | 1 | 436 |
| Arsenic (As) | 1 | 1 | 1 | 1869 |
| Cadmium (Cd) | 1 | 1 | 1 | 99 |
| Chromium (Cr) | 1 | 1 | 1 | 1129 |
| Cobalt (Co) | 0.96 | 1 | 0.98 | 185 |
| Copper (Cu) | 1 | 0.96 | 0.98 | 3451 |
| Iron (Fe) | 1 | 1 | 1 | 1147 |
| Lead (Pb) | 1 | 1 | 1 | 8 |
| Magnesium (Mg) | 1 | 1 | 1 | 10 |
| Manganese (Mn) | 1 | 1 | 1 | 1 |
| Mercury (Hg) | 1 | 1 | 1 | 366 |
| Nickel (Ni) | 0.94 | 1 | 0.97 | 679 |
| Silver (Ag) | 1 | 1 | 1 | 694 |
| Zinc (Zn) | 0.98 | 1 | 0.99 | 368 |

Performance of DeepMRG for each individual metal label available in 3rd fold

| Metal | Precision | Recall | F1-score | # of sequences |
|---|---|---|---|---|
| Antimony (Sb) | 1 | 1 | 1 | 15 |
| Arsenic (As) | 1 | 1 | 1 | 116 |
| Cadmium (Cd) | 1 | 1 | 1 | 230 |
| Chromium (Cr) | 1 | 1 | 1 | 433 |
| Cobalt (Co) | 1 | 1 | 1 | 1158 |
| Copper (Cu) | 1 | 1 | 1 | 4786 |
| Gold (Au) | 1 | 1 | 1 | 265 |
| Iron (Fe) | 1 | 1 | 1 | 3 |
| Lead (Pb) | 1 | 1 | 1 | 2 |
| Magnesium (Mg) | 1 | 1 | 1 | 1 |
| Manganese (Mn) | 1 | 1 | 1 | 3 |
| Mercury (Hg) | 1 | 1 | 1 | 635 |
| Molybdenum (Mo) | 1 | 1 | 1 | 564 |
| Nickel (Ni) | 1 | 0.99 | 1 | 1197 |
| Silver (Ag) | 1 | 0.55 | 0.71 | 930 |
| Tellurium (Te) | 1 | 1 | 1 | 403 |
| Tungsten (W) | 1 | 1 | 1 | 564 |
| Zinc (Zn) | 1 | 1 | 1 | 823 |

Performance of DeepMRG for each individual metal label available in 4th fold

| Metal | Precision | Recall | F1-score | # of sequences |
|---|---|---|---|---|
| Antimony (Sb) | 1 | 1 | 1 | 474 |
| Arsenic (As) | 1 | 1 | 1 | 757 |
| Cadmium (Cd) | 1 | 0.99 | 1 | 1171 |
| Chromium (Cr) | 1 | 1 | 1 | 493 |
| Cobalt (Co) | 1 | 0.98 | 0.99 | 1915 |
| Copper (Cu) | 0.98 | 1 | 0.99 | 3826 |
| Gallium (Ga) | 1 | 1 | 1 | 145 |
| Gold (Au) | 0.98 | 1 | 0.99 | 212 |
| Iron (Fe) | 1 | 1 | 1 | 2877 |
| Lead (Pb) | 1 | 1 | 1 | 1 |

| | | | | |
|---|---|---|---|---|
| Magnesium (Mg) | 1 | 1 | 1 | 1181 |
| Manganese (Mn) | 1 | 1 | 1 | 1248 |
| Mercury (Hg) | 1 | 0.97 | 0.98 | 609 |
| Molybdenum (Mo) | 1 | 1 | 1 | 1447 |
| Nickel (Ni) | 1 | 1 | 1 | 2363 |
| Selenium (Se) | 1 | 1 | 1 | 493 |
| Silver (Ag) | 1 | 0.59 | 0.74 | 665 |
| Tellurium (Te) | 1 | 1 | 1 | 1000 |
| Tungsten (W) | 1 | 1 | 1 | 1920 |
| Zinc (Zn) | 1 | 1 | 1 | 3198 |

Performance of DeepMRG for each individual metal label available in $5^{th}$ fold

| Metal | Precision | Recall | F1-score | # of sequences |
|---|---|---|---|---|
| Antimony (Sb) | 1 | 1 | 1 | 273 |
| Arsenic (As) | 1 | 1 | 1 | 1852 |
| Cadmium (Cd) | 0.98 | 1 | 0.99 | 1432 |
| Chromium (Cr) | 1 | 1 | 1 | 764 |
| Cobalt (Co) | 1 | 1 | 1 | 1296 |
| Copper (Cu) | 1 | 1 | 1 | 2450 |
| Gallium (Ga) | 1 | 0.73 | 0.85 | 127 |
| Gold (Au) | 1 | 1 | 1 | 299 |
| Iron (Fe) | 1 | 0.95 | 0.98 | 1851 |
| Lead (Pb) | 0.3 | 1 | 0.46 | 8 |
| Magnesium (Mg) | 1 | 1 | 1 | 922 |
| Manganese (Mn) | 1 | 1 | 1 | 992 |
| Mercury (Hg) | 1 | 1 | 1 | 1809 |
| Molybdenum (Mo) | 1 | 1 | 1 | 769 |
| Nickel (Ni) | 1 | 0.96 | 0.98 | 1565 |
| Selenium (Se) | 1 | 1 | 1 | 764 |
| Silver (Ag) | 0.78 | 1 | 0.88 | 496 |
| Tellurium (Te) | 1 | 1 | 1 | 854 |
| Tungsten (W) | 1 | 1 | 1 | 1186 |
| Zinc (Zn) | 1 | 0.95 | 0.98 | 2302 |
