## Supplementary material for "DeepMRG: a multi-label deep learning classifier for predicting bacterial metal resistance genes": S6 File

### S6 Detailed results of DeepMRG, AMRFinderPlus, and BLAST best hit methods under different parameters on the sequences in BacMet Predicted MRG DB partition 2

Precision, recall, and F1-score of DeepMRG, AMRFinderPlus, and BLAST best hit approaches (with different parameters) for each individual metal label available in BacMet Predicted MRG DB partition 2.

Here, BLAST best hit (x%, y, z%) refers to the best alignment hit with identity ≥ x%, e-value ≤ y, and coverage ≥ z%. The precision, recall, and F1-score values for metal categories where AMRFinderPlus does not provide predictions are denoted as N/A.

| Metal | Methods | Precision | Recall | F1-score | # of sequences |
|---|---|---|---|---|---|
| Antimony (Sb) | DeepMRG | 1 | 1 | 1 | 47 |
| | AMRFinderPlus | N/A | N/A | N/A | |
| | BLAST best hit (40%, 1e-7, 60%) | 1 | 1 | 1 | |
| | BLAST best hit (30%, 1e-7, 60%) | 1 | 1 | 1 | |
| Arsenic (As) | DeepMRG | 1 | 1 | 1 | 635 |
| | AMRFinderPlus | 0 | 0 | 0 | |
| | BLAST best hit (40%, 1e-7, 60%) | 1 | 0.87 | 0.93 | |
| | BLAST best hit (30%, 1e-7, 60%) | 1 | 1 | 1 | |
| Cadmium (Cd) | DeepMRG | 1 | 0.96 | 0.98 | 714 |
| | AMRFinderPlus | 0 | 0 | 0 | |
| | BLAST best hit (40%, 1e-7, 60%) | 1 | 0.9 | 0.95 | |
| | BLAST best hit (30%, 1e-7, 60%) | 1 | 1 | 1 | |
| Chromium (Cr) | DeepMRG | 1 | 1 | 1 | 20 |
| | AMRFinderPlus | 0 | 0 | 0 | |
| | BLAST best hit (40%, 1e-7, 60%) | 1 | 1 | 1 | |

| | | | | | |
|---|---|---|---|---|---|
| | BLAST best hit (30%, 1e-7, 60%) | 1 | 1 | 1 | |
| Cobalt (Co) | DeepMRG | 1 | 0.84 | 0.91 | 499 |
| | AMRFinderPlus | 0 | 0 | 0 | |
| | BLAST best hit (40%, 1e-7, 60%) | 0.99 | 0.75 | 0.85 | |
| | BLAST best hit (30%, 1e-7, 60%) | 0.94 | 1 | 0.97 | |
| Copper (Cu) | DeepMRG | 1 | 1 | 1 | 2699 |
| | AMRFinderPlus | 0 | 0 | 0 | |
| | BLAST best hit (40%, 1e-7, 60%) | 1 | 0.81 | 0.9 | |
| | BLAST best hit (30%, 1e-7, 60%) | 1 | 1 | 1 | |
| Iron (Fe) | DeepMRG | 1 | 1 | 1 | 230 |
| | AMRFinderPlus | N/A | N/A | N/A | |
| | BLAST best hit (40%, 1e-7, 60%) | 1 | 1 | 1 | |
| | BLAST best hit (30%, 1e-7, 60%) | 1 | 1 | 1 | |
| Manganese (Mn) | DeepMRG | 1 | 1 | 1 | 288 |
| | AMRFinderPlus | N/A | N/A | N/A | |
| | BLAST best hit (40%, 1e-7, 60%) | 1 | 1 | 1 | |
| | BLAST best hit (30%, 1e-7, 60%) | 1 | 1 | 1 | |
| Mercury (Hg) | DeepMRG | 1 | 0.98 | 0.99 | 710 |
| | AMRFinderPlus | 1 | 0.33 | 0.5 | |
| | BLAST best hit (40%, 1e-7, 60%) | 0.99 | 0.85 | 0.92 | |
| | BLAST best hit (30%, 1e-7, 60%) | 0.99 | 0.93 | 0.96 | |
| Molybdenum (Mo) | DeepMRG | 1 | 1 | 1 | 571 |
| | AMRFinderPlus | N/A | N/A | N/A | |
| | BLAST best hit (40%, 1e-7, 60%) | 1 | 0.65 | 0.79 | |

| | | | | | |
|---|---|---|---|---|---|
| | BLAST best hit (30%, 1e-7, 60%) | 1 | 1 | 1 | |
| Nickel (Ni) | DeepMRG | 1 | 1 | 1 | 290 |
| | AMRFinderPlus | 0 | 0 | 0 | |
| | BLAST best hit (40%, 1e-7, 60%) | 1 | 0.99 | 1 | |
| | BLAST best hit (30%, 1e-7, 60%) | 1 | 1 | 1 | |
| Silver (Ag) | DeepMRG | 0.37 | 0.95 | 0.53 | 39 |
| | AMRFinderPlus | 0 | 0 | 0 | |
| | BLAST best hit (40%, 1e-7, 60%) | 0.95 | 0.97 | 0.96 | |
| | BLAST best hit (30%, 1e-7, 60%) | 0.95 | 0.97 | 0.96 | |
| Tellurium (Te) | DeepMRG | 1 | 1 | 1 | 675 |
| | AMRFinderPlus | 0 | 0 | 0 | |
| | BLAST best hit (40%, 1e-7, 60%) | 1 | 0.37 | 0.54 | |
| | BLAST best hit (30%, 1e-7, 60%) | 1 | 0.98 | 0.99 | |
| Tungsten (W) | DeepMRG | 1 | 0.99 | 1 | 642 |
| | AMRFinderPlus | N/A | N/A | N/A | |
| | BLAST best hit (40%, 1e-7, 60%) | 1 | 0.69 | 0.81 | |
| | BLAST best hit (30%, 1e-7, 60%) | 1 | 1 | 1 | |
| Zinc (Zn) | DeepMRG | 1 | 0.98 | 0.99 | 114 |
| | AMRFinderPlus | 0 | 0 | 0 | |
| | BLAST best hit (40%, 1e-7, 60%) | 1 | 0.73 | 0.84 | |
| | BLAST best hit (30%, 1e-7, 60%) | 1 | 1 | 1 | |
