## Supplementary material for "DeepMRG: a multi-label deep learning classifier for predicting bacterial metal resistance genes": S7 File

### S7 BLAST best hit sequence identity of *Cupriavidus* strain STM 6070 heavy metal resistance (HMR) genes against the BacMet experimentally confirmed MRG database

In the following figure, each dot corresponds to the best hit of each STM 6070 HMR gene with BacMet EXP MRGs where size depicts the alignment coverage.

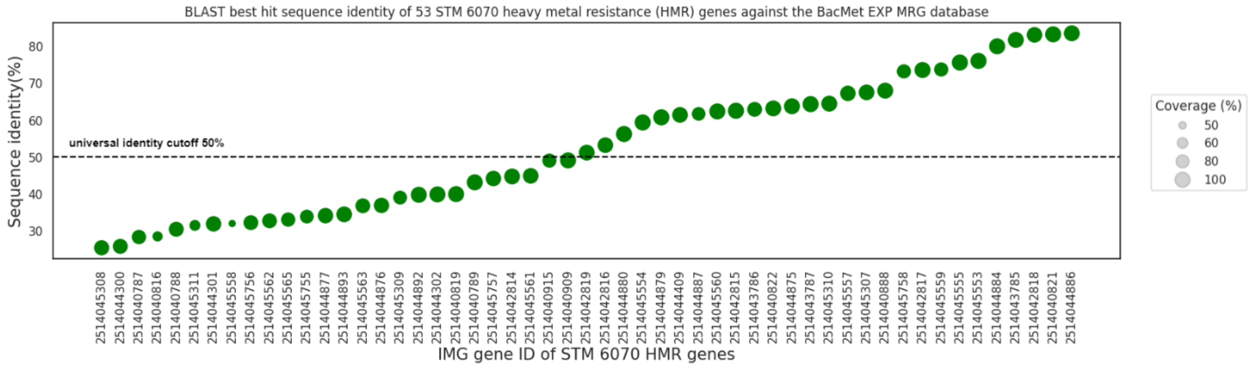


**Integrated Microbial Genomes (IMG) (https://img.jgi.doe.gov/cgi-bin/m/main.cgi?section=GeneSearch&page=searchForm)**
