## Supplementary material for "DeepMRG: a multi-label deep learning classifier for predicting bacterial metal resistance genes": S8 File

### S8 Detailed results of DeepMRG, AMRFinderPlus-50%, and BLAST-50% on the independent set of *Cupriavidus* strain STM 6070 heavy metal resistance (HMR) genes

**Table 1**: Precision, recall, and F1-score of DeepMRG, AMRFinderPlus-50%, and BLAST-50% for each individual metal label available in the independent dataset containing *Cupriavidus* strain STM 6070 HMR genes. Here, BLAST best hit (x%, y, z%) refers to the best alignment hit with identity ≥ x%, e-value ≤ y, and coverage ≥ z%.

| Metal | Methods | Precision | Recall | F1-score | # of sequences |
|---|---|---|---|---|---|
| Arsenic (As) | DeepMRG | 1 | 0.73 | 0.84 | 11 |
| | AMRFinderPlus-50% | 1 | 0.18 | 0.31 | |
| | BLAST best hit (50%, 1e-7, 60%) | 1 | 0.27 | 0.43 | |
| Cadmium (Cd) | DeepMRG | 1 | 0.57 | 0.73 | 21 |
| | AMRFinderPlus-50% | 0 | 0 | 0 | |
| | BLAST best hit (50%, 1e-7, 60%) | 1 | 0.38 | 0.55 | |
| Chromium (Cr) | DeepMRG | 1 | 1 | 1 | 5 |
| | AMRFinderPlus-50% | 0 | 0 | 0 | |
| | BLAST best hit (50%, 1e-7, 60%) | 1 | 1 | 1 | |
| Cobalt (Co) | DeepMRG | 1 | 0.50 | 0.67 | 20 |
| | AMRFinderPlus-50% | 0 | 0 | 0 | |
| | BLAST best hit (50%, 1e-7, 60%) | 0.89 | 0.40 | 0.55 | |
| Copper (Cu) | DeepMRG | 1 | 0.60 | 0.75 | 15 |
| | AMRFinderPlus-50% | 1 | 0.33 | 0.50 | |
| | BLAST best hit (50%, 1e-7, 60%) | 1 | 0.47 | 0.64 | |
| Nickel (Ni) | DeepMRG | 0.38 | 0.75 | 0.50 | 4 |
| | AMRFinderPlus-50% | 0.67 | 0.50 | 0.57 | |
| | BLAST best hit (50%, 1e-7, 60%) | 1 | 0.50 | 0.67 | |
| Silver (Ag) | DeepMRG | 0.67 | 0.29 | 0.40 | 7 |
| | AMRFinderPlus-50% | 0.50 | 0.14 | 0.22 | |
| | BLAST best hit (50%, 1e-7, 60%) | 1 | 0.14 | 0.25 | |

| | | | | | |
|---|---|---|---|---|---|
| Zinc (Zn) | DeepMRG | 1 | 0.81 | 0.89 | 21 |
| | AMRFinderPlus-50% | 0 | 0 | 0 | |
| | BLAST best hit (50%, 1e-7, 60%) | 1 | 0.38 | 0.55 | |

**Table 2**: Performance of DeepMRG on the independent dataset with the initial DIAMOND alignment parameters: sequence identity ≥ 20%, e-value ≤ 1e-3, and alignment coverage ≥ 40%.

| Method | Macro-average F1-score | Weighted-average F1-score | Samples-average F1-score |
|---|---|---|---|
| DeepMRG | 0.76 | 0.78 | 0.74 |

**Table 3**: Precision, recall, and F1-score of DeepMRG (with the initial DIAMOND alignment parameters: sequence identity ≥ 20%, e-value ≤ 1e-3, and alignment coverage ≥ 40%) for each individual metal label available in the independent dataset.

| Metal | Method | Precision | Recall | F1-score | # of sequences |
|---|---|---|---|---|---|
| Arsenic (As) | DeepMRG | 1 | 0.82 | 0.90 | 11 |
| Cadmium (Cd) | DeepMRG | 1 | 0.67 | 0.80 | 21 |
| Chromium (Cr) | DeepMRG | 1 | 1 | 1 | 5 |
| Cobalt (Co) | DeepMRG | 1 | 0.55 | 0.71 | 20 |
| Copper (Cu) | DeepMRG | 1 | 0.53 | 0.70 | 15 |
| Nickel (Ni) | DeepMRG | 0.38 | 0.75 | 0.50 | 4 |
| Silver (Ag) | DeepMRG | 0.75 | 0.43 | 0.55 | 7 |
| Zinc (Zn) | DeepMRG | 1 | 0.81 | 0.89 | 21 |
