## Supplementary material for "DeepMRG: a multi-label deep learning classifier for predicting bacterial metal resistance genes": S9 File

### S8 GO terms of the bacterial housekeeping genes that were used to create the negative microbial dataset

aerobic respiration [GO:0009060]

carbon catabolite repression of transcription [GO:0045013]

transcription-coupled nucleotide-excision repair [GO:0006283]

transcription-coupled nucleotide-excision repair, DNA damage recognition [GO:0000716]

nucleotide-excision repair, DNA damage recognition [GO:0000715]

nucleotide-excision repair, preincision complex assembly [GO:0006294]

base-excision repair, DNA ligation [GO:0006288]

nucleotide-excision repair [GO:0006289]

base-excision repair, AP site formation via deaminated base removal [GO:0097510]

nucleotide-excision repair, DNA incision [GO:0033683]

base-excision repair [GO:0006284]

nucleotide-excision repair, DNA duplex unwinding [GO:0000717]

base-excision repair, AP site formation [GO:0006285]

canonical glycolysis [GO:0061621]

acyl-CoA metabolic process [GO:0006637]

one-carbon metabolic process [GO:0006730]

lipoprotein metabolic process [GO:0042157]

regulation of RNA metabolic process [GO:0051252]

hypoxanthine metabolic process [GO:0046100]

amine metabolic process [GO:0009308]

inositol metabolic process [GO:0006020]

UDP-glucose metabolic process [GO:0006011]

ribose phosphate metabolic process [GO:0019693]

sulfur amino acid metabolic process [GO:0000096]

3'-phosphoadenosine 5'-phosphosulfate metabolic process [GO:0050427]

regulation of glutamine family amino acid metabolic process [GO:0000820]

dodecyl sulfate metabolic process [GO:0018909]

fructose 1,6-bisphosphate metabolic process [GO:0030388]

nitrate metabolic process [GO:0042126]

glycerol to glycerone phosphate metabolic process [GO:0061610]

glutathione metabolic process [GO:0006749]

glucose 6-phosphate metabolic process [GO:0051156]

GDP-mannose metabolic process [GO:0019673]

carbohydrate derivative metabolic process [GO:1901135]

regulation of cellular organohalogen metabolic process [GO:0090347]

biotin metabolic process [GO:0006768]

succinate metabolic process [GO:0006105]

aminophosphonate metabolic process [GO:0033051]

N-acetylneuraminate metabolic process [GO:0006054]

siderophore metabolic process [GO:0009237]

nucleobase-containing compound metabolic process [GO:0006139]

dihydrofolate metabolic process [GO:0046452]

folic acid-containing compound metabolic process [GO:0006760]

glutamine metabolic process [GO:0006541]

L-lyxose metabolic process [GO:0019324]

IMP metabolic process [GO:0046040]

galactitol metabolic process [GO:0019402]

shikimate metabolic process [GO:0019632]

D-amino acid metabolic process [GO:0046416]

peptidoglycan metabolic process [GO:0000270]

nitroglycerin metabolic process [GO:0018937]

nucleotide metabolic process [GO:0009117]

hydrogen peroxide metabolic process [GO:0042743]

S-methylmethionine metabolic process [GO:0033477]

regulation of nitrogen compound metabolic process [GO:0051171]

cellular aromatic compound metabolic process [GO:0006725]

positive regulation of secondary metabolite biosynthetic process [GO:1900378]

regulation of tryptophan metabolic process [GO:0090357]

positive regulation of cellular carbohydrate metabolic process [GO:0010676]

nucleoside phosphate metabolic process [GO:0006753]

L-ascorbic acid metabolic process [GO:0019852]

succinyl-CoA metabolic process [GO:0006104]

thiamine metabolic process [GO:0006772]

fucose metabolic process [GO:0006004]

amino sugar metabolic process [GO:0006040]

coenzyme A metabolic process [GO:0015936]

purine nucleotide metabolic process [GO:0006163]

D-xylose metabolic process [GO:0042732]

sulfur compound metabolic process [GO:0006790]

heme metabolic process [GO:0042168]

glutamate metabolic process [GO:0006536]

glycerol-3-phosphate metabolic process [GO:0006072]

glucose metabolic process [GO:0006006]

malate metabolic process [GO:0006108]

positive regulation of carbohydrate metabolic process [GO:0045913]

alpha-amino acid metabolic process [GO:1901605]

phosphatidylglycerol metabolic process [GO:0046471]

lipoate metabolic process [GO:0009106]

thymidine metabolic process [GO:0046104]

secondary metabolite biosynthetic process [GO:0044550]

pyrimidine nucleobase metabolic process [GO:0006206]

UDP-N-acetylglucosamine metabolic process [GO:0006047]

trehalose metabolism in response to cold stress [GO:0070415]

pyrimidine nucleotide metabolic process [GO:0006220]

fatty acid metabolic process [GO:0006631]

fumarate metabolic process [GO:0006106]

nucleobase-containing small molecule metabolic process [GO:0055086]

rhamnose metabolic process [GO:0019299]

peptide metabolic process [GO:0006518]

carboxylic acid metabolic process [GO:0019752]

acetate metabolic process [GO:0006083]

NADP metabolic process [GO:0006739]

N-acetylglucosamine metabolic process [GO:0006044]

diacetylchitobiose metabolic process [GO:0052778]

carbohydrate metabolic process [GO:0005975]

D-gluconate metabolic process [GO:0019521]

cellular carbohydrate metabolic process [GO:0044262]

selenocysteine metabolic process [GO:0016259]

acetyl-CoA metabolic process [GO:0006084]

argininosuccinate metabolic process [GO:0000053]

galactarate metabolic process [GO:0019580]

guanosine tetraphosphate metabolic process [GO:0015969]

generation of precursor metabolites and energy [GO:0006091]

NADH metabolic process [GO:0006734]

negative regulation of cellular carbohydrate metabolic process [GO:0010677]

aspartate metabolic process [GO:0006531]

nitrogen compound metabolic process [GO:0006807]

colanic acid metabolic process [GO:0046377]

organic phosphonate metabolic process [GO:0019634]

superoxide metabolic process [GO:0006801]

pyrimidine nucleoside metabolic process [GO:0006213]

L-fucose metabolic process [GO:0042354]

nucleoside monophosphate metabolic process [GO:0009123]

glycerol metabolic process [GO:0006071]

fructose metabolic process [GO:0006000]

metabolic process [GO:0008152]

toxic metabolite repair [GO:0110052]

propionate metabolic process, methylcitrate cycle [GO:0019679]

threonine metabolic process [GO:0006566]

negative regulation of carbohydrate metabolic process [GO:0045912]

tetrahydrofolate metabolic process [GO:0046653]

phosphatidylcholine metabolic process [GO:0046470]

purine nucleobase metabolic process [GO:0006144]

phosphorus metabolic process [GO:0006793]

citrate metabolic process [GO:0006101]

lipid metabolic process [GO:0006629]

D-glucarate metabolic process [GO:0042836]

trehalose metabolic process [GO:0005991]

D-serine metabolic process [GO:0070178]

pyridoxal metabolic process [GO:0042817]

nucleoside diphosphate metabolic process [GO:0009132]

fructose 6-phosphate metabolic process [GO:0006002]

lipopolysaccharide metabolic process [GO:0008653]

N-acetylmannosamine metabolic process [GO:0006051]

tartrate metabolic process [GO:1901275]

2-oxoglutarate metabolic process [GO:0006103]

regulation of fatty acid metabolic process [GO:0019217]

xenobiotic metabolic process [GO:0006805]

negative regulation of phosphate metabolic process [GO:0045936]

D-ribose metabolic process [GO:0006014]

riboflavin metabolic process [GO:0006771]

ferredoxin metabolic process [GO:0006124]

oxaloacetate metabolic process [GO:0006107]

cytosine metabolic process [GO:0019858]

positive regulation of tryptophan metabolic process [GO:0090358]

NAD metabolic process [GO:0019674]

primary metabolic process [GO:0044238]

protoporphyrinogen IX metabolic process [GO:0046501]

cellular metabolic compound salvage [GO:0043094]

carnitine metabolic process [GO:0009437]

propanediol metabolic process [GO:0051143]

regulation of secondary metabolite biosynthetic process [GO:1900376]

glyoxylate metabolic process [GO:0046487]

mannitol metabolic process [GO:0019594]

short-chain fatty acid metabolic process [GO:0046459]

formaldehyde metabolic process [GO:0046292]

dimethyl sulfoxide metabolic process [GO:0018907]

metabolite repair [GO:0110051]

glycerophospholipid metabolic process [GO:0006650]

pyruvate metabolic process [GO:0006090]

phosphate-containing compound metabolic process [GO:0006796]

galactose metabolic process [GO:0006012]

purine nucleoside metabolic process [GO:0042278]

ribonucleoside diphosphate metabolic process [GO:0009185]

cellular amino acid metabolic process [GO:0006520]

mannose metabolic process [GO:0006013]

cellular lipid metabolic process [GO:0044255]

nicotinamide nucleotide metabolic process [GO:0046496]

homocysteine metabolic process [GO:0050667]

folic acid metabolic process [GO:0046655]

tRNA metabolic process [GO:0006399]

chorismate metabolic process [GO:0046417]

| |
|---|
| L-serine metabolic process [GO:0006563] |
| guanine metabolic process [GO:0046098] |
| nucleotide-sugar metabolic process [GO:0009225] |
| nucleoside metabolic process [GO:0009116] |
| arginine metabolic process [GO:0006525] |
| spermidine metabolic process [GO:0008216] |
| cellular biogenic amine metabolic process [GO:0006576] |
| DNA metabolic process [GO:0006259] |
| positive regulation of isoprenoid metabolic process [GO:0045828] |
| pentose-phosphate shunt [GO:0006098] |
| pentose-phosphate shunt, oxidative branch [GO:0009051] |
| pentose-phosphate shunt, non-oxidative branch [GO:0009052] |
